# Plasma Exosomes in Insulin Resistant Obesity Exacerbate Progression of Triple Negative Breast Cancer

**DOI:** 10.1101/2024.10.10.617639

**Authors:** Pablo Llevenes, Andrew Chen, Matthew Lawton, Alejandro N. Rondón-Ortiz, Yuhan Qiu, Michael Seen, Stefano Monti, Gerald V. Denis

## Abstract

Breast cancer, the most common cancer among women worldwide, continues to pose significant public health challenges. Among the subtypes of breast cancer, triple-negative breast cancer (TNBC) is particularly aggressive and difficult to treat due to the absence of receptors for estrogen, progesterone, or human epidermal growth factor receptor 2, rendering TNBC refractory to conventional targeted therapies. Emerging research underscores the exacerbating role of metabolic disorders, such as type 2 diabetes and obesity, on TNBC aggressiveness. Here, we investigate the critical cellular and molecular factors underlying this link. We explore the pivotal role of circulating plasma exosomes in modulating the tumor microenvironment and enhancing TNBC aggressiveness. We find that plasma exosomes from diet-induced obesity mice induce epithelial-mesenchymal transition features in TNBC cells, leading to increased migration *in vitro* and enhanced metastasis *in vivo*. We build on our previous reports demonstrating that plasma exosomes from obese, diabetic patients, and exosomes from insulin-resistant 3T3-L1 adipocytes, upregulate key transcriptional signatures of epithelial-mesenchymal transition in breast cancer. Bioinformatic analysis reveals that TNBC cells exhibit higher expression and activation of proteins related to the Rho-GTPase cascade, particularly the small Ras-related protein Rac1. Our approach suggests novel therapeutic targets and exosomal biomarkers, ultimately to improve prognosis for TNBC patients with co-morbid metabolic disorders.

---

Breast cancer is the most common cancer among women worldwide and represents a major public health concern due to its high incidence and significant mortality rates. In the United States alone, approximately 297,790 new cases of invasive breast cancer were reported in women in 2023, resulting in an estimated 43,700 deaths. Additionally, around 310,720 new cases of invasive breast cancer and about 56,500 new cases of ductal carcinoma in situ are expected to be diagnosed in women in 2024, with an estimated 42,250 deaths resulting from the disease^1^.

Among the various subtypes of breast cancer, triple-negative breast cancer (TNBC) stands out due to its aggressive nature and poor prognosis. TNBC is characterized by the absence of three critical receptors: estrogen receptor (ER), progesterone receptor, and human epidermal growth factor receptor 2 (HER2), which are common targets for breast cancer therapies^2^. Consequently, TNBC lacks targeted treatment options, relying primarily on conventional cytotoxic therapies. This limitation contributes to the subtype’s high recurrence rates and significantly lower survival rates compared to other forms of breast cancer^3^. While other types of invasive breast cancer, such as localized ER-positive and HER2-positive, have a 5-year relative survival rate of around 99% and 93%, respectively, the 5-year relative survival rate for TNBC is approximately 91%. However, this rate drops dramatically to about 12% for distant metastases^4^. This stark decrease underscores the urgent need for more effective treatment strategies for metastatic TNBC.

Recent years have seen a growing interest in the intersection of TNBC with metabolic disorders, such as type 2 diabetes (T2D) and obesity, as these conditions are linked to heightened cancer aggressiveness and poorer patient outcomes^5^. Obesity, as a chronic inflammatory condition, disrupts tissue insulin sensitivity and function, leading to the development of T2D and adipocyte insulin resistance (IR)^6^. This systemic inflammation causes metabolic abnormalities that release various pro-inflammatory factors, such as cytokines interleukin (IL)-6, tumor necrosis factor (TNF)-α, and leptin, into the plasma^7,8^. The inflammatory milieu created by obesity promotes cancer development and progression through the interactions of these circulating factors with the tumor and its microenvironment^9,10^.

Emerging research has increasingly focused on the role of exosomes in cancer aggressiveness. Exosomes are small extracellular vesicles (∼100nm) found in blood and other fluids, containing proteins, lipids, and nucleic acids that reflect the content of their cells of origin. These vesicles facilitate intercellular communication and can reprogram recipient cells^11^. Depending on the metabolic state of the originating cell, exosomes can create a synergistic environment that promotes processes in malignant cells, such as epithelial-to-mesenchymal transition (EMT) and cancer stem cell (CSC) formation, thereby contributing to a more aggressive tumor phenotype^12,13^.

Our research builds upon this foundation by exploring a novel aspect of the interaction between metabolic disorders and TNBC aggressiveness. Our preliminary studies have demonstrated that exosomes from IR adipocytes promote significant EMT and tumor progression in breast cancer models, compared to insulin-sensitive controls^14^. We have also observed that exosomes from T2D patient plasma contain miRNAs that enhance tumor cell invasiveness and metastatic potential compared to non-diabetic controls^15,16^. Given the substantial impact of metabolic diseases on TNBC outcomes, our research aims to delve deeper into the molecular mechanisms underlying this interaction.

We hypothesize that exosomes derived from adipocytes and plasma play a pivotal role in modulating the tumor microenvironment to favor tumor aggressiveness and metastasis. We used murine models in our approach to test this hypothesis. Exosomes from adipocytes and plasma of mice maintained on a high-fat diet (HFD) for 12 weeks carry unique biomolecules that dramatically alter the behavior of TNBC cells, promoting processes such as EMT and enhanced metastatic potential, unlike matched exosomes from low-fat diet (LFD) controls.

Our study offers a novel contribution by investigating the functional impact of these exosomes across different metabolic states, with the goal of identifying new therapeutic targets and biomarkers. This research has the potential to reveal previously unrecognized mechanisms by which metabolic dysregulation drives TNBC progression, ultimately informing more effective strategies for managing this aggressive breast cancer subtype in patients with co-morbid metabolic disorders.

## Materials and Methods

### Cell culture

Murine E0771-GFP (green fluorescence protein) cells as a syngeneic TNBC model for C57BL/6J mice were a gift of Dr. Timothy Padera. The cells were cultured in Dulbecco’s Modified Eagle Medium (DMEM) medium with 4.5 g/L glucose and L-glutamine, without sodium pyruvate, supplemented with 10% fetal bovine serum (FBS), 100 units/mL penicillin and 10 μg/mL streptomycin (Corning), and incubated at 37°C with 100% humidity and 5% CO_2_. Cell lines were tested monthly for mycoplasma using a Mycoplasma Detection Kit (Invivogen) and used within 10 passages.

### Animals

Six-week-old female C57BL/6J mice were obtained from Jackson Laboratory and acclimated at the Boston University Animal Facility for one week. Mice were fed *ad libitum* a lard-based high-fat diet (HFD, 60% of calories from fat, Research Diets D12492i) or a matched low-fat diet (LFD, 10% fat, Research Diets D12450Ji) for 12 weeks, ensuring obesity in the HFD group. All animal experiments followed the approved procedures of the Institutional Animal Care and Use Committee (IACUC) of Boston University Medical Center. For all consequent experiments, E0771-GFP cells were treated with exosomes on Day 0 and allowed to proliferate for 72 hours up to a confluence of 70%.

### Animal handling and euthanasia

All euthanasia procedures were conducted in accordance with the IACUC guidelines at Boston University Medical Center to ensure humane treatment and minimize distress. Euthanasia was carried out using an isoflurane inhalation system followed by cervical dislocation for confirmation of death. Mice were placed in an induction chamber connected to an isoflurane delivery system. This system included an inlet tube that delivered a controlled mixture of isoflurane and oxygen into the chamber and an outlet tube that vented residual gas to a scavenging system for safe disposal. The isoflurane intensity was set to a level suitable for inducing deep anesthesia in rodents. While oxygen levels were kept minimal, a low oxygen flow was maintained to ensure effective delivery of isoflurane. The animals were closely monitored during the procedure to confirm deep anesthesia, indicated by complete loss of toe pinch reflexes and slowed respiration. Once unconsciousness was verified, cervical dislocation was performed to confirm death and ensure ethical and definitive euthanasia. This combination of isoflurane inhalation and cervical dislocation ensured both humane induction of unconsciousness and reliable confirmation of death.

### Glucose tolerance test, insulin tolerance tests, weight curves and body mass index

Glucose tolerance tests (GTTs) and insulin tolerance tests (ITTs) were performed on adult female mice (18 weeks old, with 12 weeks on diet: N = 5 HFD and 5 matching control LFD). For GTT, after a 6-hour fast, mice were administered a glucose solution via oral gavage at a dose of 2 g/kg body weight. Blood glucose levels were measured at baseline (time = 0) and at 15, 30, 60 and 120 minutes post-glucose administration using a glucometer (Abbott, AlphaTrak2) with blood samples collected from the tail vein. For ITT, human recombinant insulin (Gibco; 0.75 U/kg body weight) from a stock solution of 4 mg/mL was administered intraperitoneally after a 6-hour fast. Blood was collected from the tail vein immediately before insulin injection (time = 0) and at 15, 30, 60, and 120 minutes after injection. Blood glucose levels were then plotted to generate glucose and insulin tolerance curves, and the area under the curve (AUC) was calculated to compare glucose and insulin tolerance between the obese and control groups. Throughout the tests, mice were observed for any signs of hypoglycemia or adverse reactions.

The weight of the mice was measured weekly for both the HFD and LFD groups throughout the 12-week diet period. At the end of the 12 weeks, body mass index (BMI) was calculated to confirm the obese status of the HFD group compared to the LFD group. BMI was determined using the formula: BMI = weight (g) / length (cm)^2. This calculation ensured an accurate assessment of obesity status in the HFD group relative to the control LFD group.

### Isolation of primary adipocytes and blood plasma from mouse

After completing the 12-week diet, mice were euthanized, and their visceral adipose tissue depots (perigonadal) were excised. The tissue was minced in cold phosphate buffered saline (PBS) pH 7.4 in a tissue culture dish and filtered through a 250-micron mesh. The minced tissue was then transferred to a 50 mL Falcon tube containing liberase (Sigma Aldrich) digestion buffer (10 mg liberase, 300 μL dH_2_O, 500 μL of 1M HEPES, pH 7.4) in a total volume of 10 mL DMEM without FBS. Tubes were placed in a shaker at 37°C, with oscillation at 120 revolutions per minute (RPM) for at least one hour. The duration of shaking was adjusted based on the initial size of the depot to ensure thorough digestion. After digestion, the mixture was filtered through a 250-micron mesh and transferred to a new tube. To neutralize the digestion, 5 mL of FBS was added. The sample was then centrifuged at 500 rcf for 5 minutes, followed by vigorous shaking and another centrifugation at 500 rcf for 5 minutes at room temperature. The layer of lipid-laden adipocytes floating at the top of the suspension was isolated and placed in 10 mL of complete DMEM in a 10 cm dish and incubated at 37°C for 24 hours to allow the exosome release into the media for collection.

Retro-orbital blood extraction was performed using capillary and collection tubes, both treated with EDTA to prevent coagulation. The initial blood volume collected after opening the wound was discarded and blood samples were kept at room temperature for less than one hour. The collected blood was first centrifuged at 250 rcf for 25 minutes at room temperature. The supernatant (plasma) was then carefully transferred to a new tube and centrifuged at 2,500 rcf for 20 minutes. The resulting plasma was collected, and the pellet was discarded. Plasma samples were stored at −80°C until further exosome purification.

### Exosome purification

Exosomes were either isolated from visceral adipose tissue or mouse plasma. For exosomes isolated from visceral adipose tissue, conditioned media (typically 15 mL) was centrifuged at 300 × g for 10 min in a 15 mL conical tube to remove cells and debris. The supernatant was then transferred to an Amicon Ultra-15 (Millipore-Sigma; REF-UFC910008) 100K centrifugal filter and centrifuged at 3,500 × g for 15 min to reach a final volume of 0.5 ml. Exosomes were then purified by size exclusion chromatography on a qEV Original column, using an automatic fraction collector (AFC, serial number: V1-0395, IZON), as previously described^15^. For exosome isolation from plasma, approximately 20 μL of plasma was processed using 35 nm qEVsingle Gen 2 columns. The columns were set with default AFC parameters. The collected fraction contained the exosomes and was diluted 1:100 (exosomes:PBS+EDTA). The size distribution and concentration of exosomes were determined before each biological experiment using the NanoSight NS300 system (Malvern Panalytical). To further confirm exosome purity, different pooled exosome fractions were tested for CD63 expression by dot blot. Equal volumes of exosome suspensions were spotted onto nitrocellulose membranes and probed with an anti-CD63 antibody. Exosome morphology was imaged using a transmission electron microscope operated at 120 kV and equipped with a NANOSPRT43 camera. Representative images were acquired at 10,000× magnification showing vesicles with characteristic size and morphology consistent with exosomes. Based on this quantitation, the ratio of exosomes to E0771 cells was 100,000:1 on Day 0 of exposure.

### PCR array

Total RNA was extracted from mouse E0771 cells using the RNeasy Plus Mini Kit (Qiagen, 74136). From each sample, 1 μg of RNA was used to prepare 20 μL of cDNA using the QuantiTect Reverse Transcription Kit (Qiagen, 205313). The Mouse RT2 Profiler™ PCR Array for Epithelial to Mesenchymal Transition (EMT) (Qiagen, PAMM-090Z) was utilized for this study. Each cDNA sample (102 μL) was mixed with 1,350 μL of RT2 SYBR Green-ROX qPCR Mastermix (Qiagen, 330522) and 1,248 μL of RNAse-free water. This brought the total volume to 2,700 μL, which was thoroughly mixed by pipetting. Using an automated pipettor, 25 μL aliquots of the reaction mix were distributed into each well of the array. Real-time PCR was conducted using the StepOne Plus PCR instrument (Applied Biosystems) following the manufacturer’s instructions. The results were analyzed and plotted using the RT2 Profiler web-based software provided by Qiagen®.

### Ingenuity pathway analysis

To predict disease functions and downstream pathways, data were analyzed using Ingenuity Pathway Analysis (IPA) software (QIAGEN®). Core analysis was first conducted, followed by comparison analysis on all experimental conditions, including replicates. This approach enabled the identification and interpretation of biological pathways, molecular networks, and functional connections relevant to the dataset.

### Small RNA Sequencing and Analysis

Exosomal miRNA was purified by phenol:chloroform extraction and ethanol precipitation. The resulting RNA was sequenced using Novogene’s small RNA sequencing service, which encompasses extraction, library preparation, and analysis. Total RNA was first isolated, followed by selection of small RNAs within the 25-200 nt range. Abclonal Small RNA Library Prep Kit for Illumina was used to capture miRNAs during the library preparation process. Small RNA fragments were ligated with adapters, reverse transcribed to cDNA, and amplified through PCR. Quality control was performed to ensure high data integrity, and sequencing was conducted on the Illumina platform with standard bioinformatic processing.

### mRNA sequencing and analysis

Total RNA was extracted from the cells using the RNeasy Plus Mini Kit (QIAGEN, Cat. No. 74136) following the manufacturer’s protocol. RNA sequencing data was processed using the nf-core rnaseq pipeline^17^, and the resulting expression matrices were filtered to exclude genes with low expression (fewer than 3 reads per million in at least 3 samples) and corrected for library size differences using the DESeq2 package^18^. Differential gene expression analysis was then performed using DESeq2 between the Control (for *in vitro* analysis LFD, and for brain metastasis analysis LB plus CB) and HFD experimental groups. A list of the genes that were used to stratify the TCGA is provided as Supplementary Table T1. We stratified TCGA breast cancer samples into molecular subtypes—Luminal A, Luminal B, HER2-enriched, and Basal-like—based on gene expression profiles. While triple-negative breast cancer (TNBC) is not explicitly annotated in TCGA, the Basal-like subtype significantly overlaps with TNBC cases. To examine the prognostic value of these DEGs in human tumors, survival analysis was conducted using a Cox proportional hazards model and visualized using the survminer package^19^ in breast cancer data from both The Cancer Genome Atlas (TCGA)^20^. The Cox model also included patient age and proliferation expression as covariates to estimate prognostic value independent of age and proliferation-related mechanisms^21^.

### Cell migration assay

Cells were initially treated with exosomes as previously described, then switched from normal medium to serum-free medium for 3 hours. After this period, cells were trypsinized, counted, and 30,000 cells were seeded into the upper wells of 24-well transwell plates (8-μm pore size, Thermo Fisher Scientific®) with 300 μL of serum-free medium. The bottom wells were filled with 500 μL of DMEM media supplemented with 10% FBS to serve as a chemoattractant. The cells were then allowed to migrate for 24 hours. After the migration period, the media was aspirated, and the transwell inserts were gently washed with PBS. The upper side of the membrane was carefully cleaned to remove non-migrated cells using a cotton swab with a gentle swirling motion, taking care not to press too hard to avoid background signals and disruption of the filter monolayer. The migrated cells on the lower side of the membrane were fixed with 500 μL of cold 100% methanol for 10 minutes at −20°C. After fixation, the methanol was aspirated, and inserts were cleaned with PBS. Once the inserts were dry, the cells were stained with 500 μL of 1% crystal violet (w/v) in 2% ethanol (v/v) for 10 minutes at room temperature. The crystal violet solution was then aspirated, and the inserts were washed three times with PBS. The inserts were allowed to air dry for 30 minutes. Images of the stained migrated cells were captured using an EVOS® XL Core digital inverted microscope. The percentage of migration was quantified using ImageJ software (National Institutes of Health, Bethesda, MD).

### Cellular symmetry analysis

Cell images were captured using an EVOS XL Core digital inverted microscope. Cell morphological parameters were analyzed using CellProfiler® software (Broad Institute, MA). The perimeter was defined as the total length of the cell’s outer boundary. Circularity was calculated using the formula 4π × (area/perimeter^2^) where a value of 1.0 denotes a perfect circle, and values approaching 0.0 indicate decreasing circularity and a more irregular shape. For cell symmetry analysis, EHT-1864 was used at 25 µM and Y-27632 at 10 µM.

### Proteomics Analysis

Samples stored at −80°C were thawed on ice and lysed using guanidine hydrochloride lysis buffer (6M guanidine hydrochloride, 100mM Tris pH 8.5, 40mM chloroacetamide, 10mM TCEP), followed by vortex mixing, heating at 95°C for 5 minutes, and sonication. Protein concentration was determined via Bradford assay. Samples were diluted to <0.75M guanidine hydrochloride using 100mM Tris-HCl (pH 8.0), digested overnight with trypsin (1:50) at 37°C, and acidified with 1% formic acid. Peptides were desalted, dried, and quantified using the Pierce peptide assay. 100 μg of peptides per sample were labeled with TMTpro, pooled, desalted, and fractionated using reversed-phase HPLC, yielding 48 fractions. For phosphopeptide enrichment, samples were resuspended in 80% acetonitrile/0.5% trifluoroacetic acid and enriched with iron-based beads (Cube Biotech). Phosphopeptides were eluted, dried, and analyzed by mass spectrometry using an Exploris 480 with FAIMS Pro. Peptides were separated by HPLC using a 120-minute gradient and analyzed in positive ion mode. Data acquisition was done in data-dependent mode, selecting the 12 most abundant ions for fragmentation. Phosphopeptides were analyzed similarly with a shorter gradient. Raw spectra were searched against the Uniprot (*Mus musculus*) database using MaxQuant. Reporter ion intensities were normalized using Loess normalization, and data were analyzed with OmicsNotebook. Differential expression was assessed using Limma, with p-values corrected via Benjamini-Hochberg. Functional enrichment analyses were performed using GSEA and EnrichR.

### Cell count

The number of cells was determined by counting triplicates for each condition. Cells were trypsinized and stained with a 1:10 trypan blue dilution (trypan blue:PBS) to distinguish viable cells from non-viable cells. The cell count was performed using a hemocytometer (Neubauer Improved) under an EVOS® XL Core digital inverted microscope.

### Clonogenic Assay

E0771-GFP cells were trypsinized and resuspended in a cold PBS solution and 100,000 were injected into the second mammary fat pad of 6-week-old C57BL/6J female mice and sacrificed four weeks post-injection. Tissues were homogenized using the previously described digestion buffer containing liberase. The resulting homogenate was filtered through cell strainer filters of 40 μm pore size, ensuring the result was a single cell suspension. Each homogenate was plated separately in 150mm culture dishes and allowed to settle. After settling, the media was gently removed and replaced to eliminate non-adherent cells, and the adherent cells were left to grow for one week to form colonies. The colonies were then analyzed using a Zeiss® Stereo Discovery V12 microscope equipped with filters for detecting GFP positive cells, along with an attached Axiocam MRC camera by Zeiss®. The images were processed using Adobe® Photoshop (Adobe Inc., San Jose, CA, USA) to create a photomerged final image, providing a comprehensive view of the whole plate. The colonies were then counted, and the cell number and the area covered by the cells on the total plate were quantified using ImageJ® software (National Institutes of Health, Bethesda, MD, USA). Primary tumors were carefully excised and measured. The dimensions of each tumor were recorded in three orthogonal directions (length, width, and height) using a digital caliper. Subsequently, the tumors were weighed using an analytical balance to obtain their mass.

### Wound healing assay

Cells were seeded for a 3-day exosome treatment in 12-well plates and allowed to reach a confluence of 90%, at which point the surface of each well was scratched using a sterile 10-µL pipette tip. The cells were then incubated overnight with increasing concentrations of the Rac1 inhibitor EHT 1864 (Cat. No. 387210R, Tocris®). EHT 1864 was prepared in a series of concentrations: 5, 10, 20, 25, 30, 40, and 50 µM. The wound closure was monitored and imaged at 0 and 24 hours post-treatment using an EVOS® XL Core digital inverted microscope with a digital camera. The wound closure was measured using ImageJ® software, and the percentage of wound closure was calculated using the formula: Percentage of Wound Closure = [(Wound Area at 0 hours -Wound Area at 24 hours) / Wound Area at 0 hours] × 100.

### Phalloidin Staining

Cells were fixed in 10% formalin for 20 minutes at room temperature, followed by three washes with PBS, then samples were permeabilized with 0.1% Triton X-100 in PBS for 20 minutes at room temperature. Cells were subsequently blocked with 1% bovine serum albumin in PBS for 30 minutes on a rocking platform at room temperature. After blocking, cells were incubated with Alexa Fluor™ 568 Phalloidin (Thermo Fisher Scientific, Cat# A12380; at 1:400 dilution in 1% BSA in PBS) for 60 minutes at room temperature on a rocking platform. After antibody incubation, samples were washed three times with PBS and counterstained with 2 μg/mL DAPI solution in PBS for 10 minutes on a rocking platform. Samples were washed three additional times with PBS and air-dried for 30 minutes prior to imaging. Imaging was performed using a Nikon Deconvolution Wide-Field Epifluorescence System. Images were analyzed using ImageJ software.

### Rac1 Activation Assay

Rac1 activation was measured using the Rac1 Activation Assay Biochem Kit™ (Cytoskeleton, Inc., Cat# BK035) following the manufacturer’s instructions. E0771 cells were treated for 72 hours with plasma-derived exosomes isolated from either HFD-fed or LFD-fed mice, or with HFD-derived exosomes in combination with the Rac1 inhibitor EHT 1864 (25 µM). Following treatment, cells were serum-starved for 3 hours, placed on ice, washed twice with ice-cold PBS, and lysed directly on the culture dishes using the provided ice-cold Cell Lysis Buffer supplemented with a 1X protease inhibitor cocktail. Cells were scraped and lysates were clarified by centrifugation at 10,000 × g for 1 minute at 4°C. Protein concentration was determined using the BCA assay (Thermo Fisher Scientific), and samples were equalized across conditions.

300 µg of total protein per sample was incubated with 10 µg of PAK-PBD-conjugated agarose beads for 1 hour at 4°C on a rotator. Beads were pelleted by centrifugation, washed with Wash Buffer, and resuspended in 2X Laemmli sample buffer. Samples were boiled for 2 minutes and subjected to SDS-PAGE, followed by western blotting. Membranes were probed with the anti-Rac1 monoclonal antibody provided in the kit (1:500 dilution), and detection was performed using an HRP-conjugated secondary antibody. Band intensity for activated Rac1 (GTP-bound) was quantified using ImageJ.

### Immunoblotting

Immunoblotting was performed to assess total Rac protein levels in E0771 cells following plasma-derived exosome treatment. Cells were washed twice with ice-cold PBS and lysed directly on the culture plates using RIPA buffer supplemented with 0.2% SDS and a protease inhibitor cocktail. Adherent cells were detached using cell scrapers. Lysates were clarified by centrifugation at 10,000 × g for 10 minutes at 4°C, and the supernatants were collected. Protein concentrations were determined using a BCA assay kit (Thermo Fisher Scientific).

Equal amounts of protein (40 µg per lane) were separated on SDS-PAGE gels and transferred onto nitrocellulose membranes. Membranes were blocked with 5% non-fat milk in Tris-buffered saline with 0.1% Tween-20 (TBS-T) for 30 minutes at room temperature. Primary antibody incubation was performed overnight at 4°C using Rac1/2/3 antibody (Cell Signaling Technology, Cat# 2465, 1:500 dilution) and β-Actin (13E5) Rabbit mAb (Cell Signaling Technology, Cat# 4970, 1:1000 dilution) as a loading control. After washing, membranes were incubated with an HRP-conjugated anti-rabbit secondary antibody (Cell Signaling Technology, Cat# 7074, 1:2000 dilution) for 1 hour at room temperature. Protein bands were visualized by enhanced chemiluminescence and imaged using a Bio-Rad ChemiDoc XRS+ chemiluminescence imaging system.

### Statistical Analyses

Statistical analyses were performed using GraphPad Prism software. Depending on the data distribution, either parametric (Student’s t-test or ANOVA) or non-parametric tests (Mann-Whitney or Kruskal-Wallis) were applied as indicated. The following symbols were used to denote the level of statistical significance: ns, p > 0.05; *, p < 0.05; **, p < 0.01; ***, p < 0.001; ****, p<0.0001. For parametric data, Student’s t-tests were used to compare two groups, and ANOVA was used for multiple group comparisons. For non-parametric data, equivalent non-parametric tests such as the Mann-Whitney U test for two groups or the Kruskal-Wallis test for multiple groups were utilized. All results are presented as mean ± standard deviation (SD) unless otherwise indicated.

## RESULTS

### Establishment of Obesity and Insulin Resistance in HFD Mice

To evaluate the effects of HFD on exosome production, six-week-old female C57BL/6J mice were fed either a lard-based high-fat diet (60% of calories from fat) or a matched low-fat diet (10% of calories from fat). After 12 weeks on diet, the HFD group developed obesity, glucose intolerance and insulin resistance, confirmed by significant weight gain and impaired glucose metabolism, as shown in Figure 1 (A-G). To confirm the obesity in the model, weekly measurements over 12 weeks demonstrated a statistically significant weight increase (Figure 1A and B), that was further confirmed by measuring the BMI at the 12^th^ week (Figure 1C), observing a BMI greater than 4 kg/m^2^ (obese) for the HFD group. To confirm the impaired glucose metabolism, oral glucose tolerance tests (oGTT) and insulin tolerance tests (ITT), were carried out (Figure 1D-G). Showing impaired glucose metabolism in the HFD group compared to LFD. These results confirmed the successful establishment of the obesity model, evidenced by significant physiological changes, including weight gain, insulin resistance, and impaired glucose metabolism.

**Figure 1.**
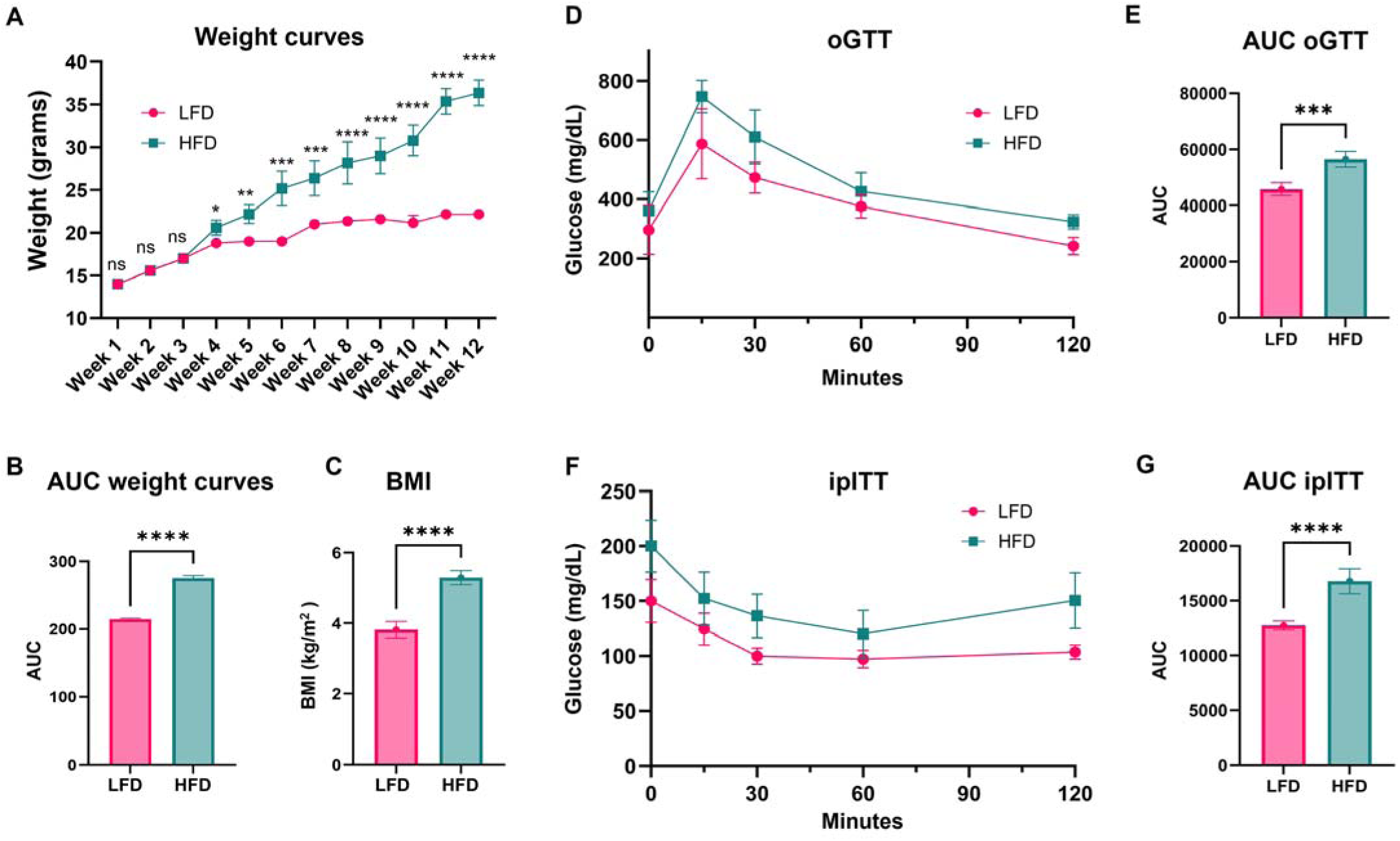
Validation of the obese mouse model. Measurements were taken after 12 wk of feeding *ad libitum* either HFD or LFD to 6-wk-old female C57BL/6J mice. (A) Weight curves over time. (B) Area under the curve (AUC) analysis of weight. (C) Body mass index (BMI, kg/m²) after 12 wk. (D) oGTT curves. (E) AUC analysis of oGTT. (F) ITT curves. (G) AUC analysis of ITT. LFD, Low-fat diet; HFD, High-fat diet; oGTT, oral glucose tolerance test; ITT, insulin tolerance test. N=5. Data were analyzed by multiple t-tests; statistical significance was ns (not significant), *p < 0.05, **p < 0.01, ***p < 0.001, ****p < 0.0001. All data are shown as mean ± SD.

After the establishment of obesity and systemic insulin resistance in the HFD group, but not in the LFD group, we proceeded with exosomal extraction. Blood was collected via retro-orbital extraction for plasma-derived exosomes, and visceral perigonadal tissue was harvested for adipocyte-derived exosome isolation. No differences were found in diameter and abundance between LFD and HFD exosomes. Furthermore, we confirmed that 20 μL of plasma yielded approximately 6 to 7 billion exosomes, providing sufficient material for our functional assays. This yield was consistent across both HFD and LFD groups. (Supplementary Figure S1A-B). The ability to obtain robust exosome quantities from minimal plasma volumes represents a strength of our approach, particularly for potential translational applications. To further confirm vesicle identity, Transmission Electron Microscopy was performed on purified exosomes showing consistent diameters with those found in the nanoparticle tracking analysis plus presence of CD63 positive vesicles of isolated extracellular vesicle fractions on chromatography (Supplementary Figure S1C-D).

### Differential Gene Expression and Morphological Changes Induced by Adipocyte-Derived Exosomes

Using exosomes from visceral fat, an EMT-array revealed differential gene expression after a three-day treatment (Figure 2A and B) (Supplementary Figure S2A, S2B). Notably, Snai1 and Ctnnb1 were significantly upregulated in HFD-treated cells compared to LFD-treated cells (Supplementary Figure S2C), as demonstrated in a volcano plot (Figure 2C). Enrichment analysis using Ingenuity Pathway Analysis (IPA) software identified the most enriched pathways as those related to cytoskeleton organization, cell survival and movement. These findings were visualized through bubble and bar charts (Figure 2D). This result prompted further analysis of cell morphology, which revealed increased perimeter and decreased circularity in HFD-treated cells (Figure 2E). These morphological changes are indicative of cytoskeletal restructuring consistent with a pro-EMT phenotype. Despite these significant molecular and morphological changes, migration assays did not show any differences in migratory capacity between HFD-treated and LFD-treated groups (Figure 2F). Similarly, cell viability assays revealed no significant differences in cell survival between the two groups (Figure 2G). These results suggest that while HFD-derived adipocyte exosomes promote molecular and structural changes associated with EMT, additional factors present in plasma exosomes account for migration differences.

**Figure 2.**
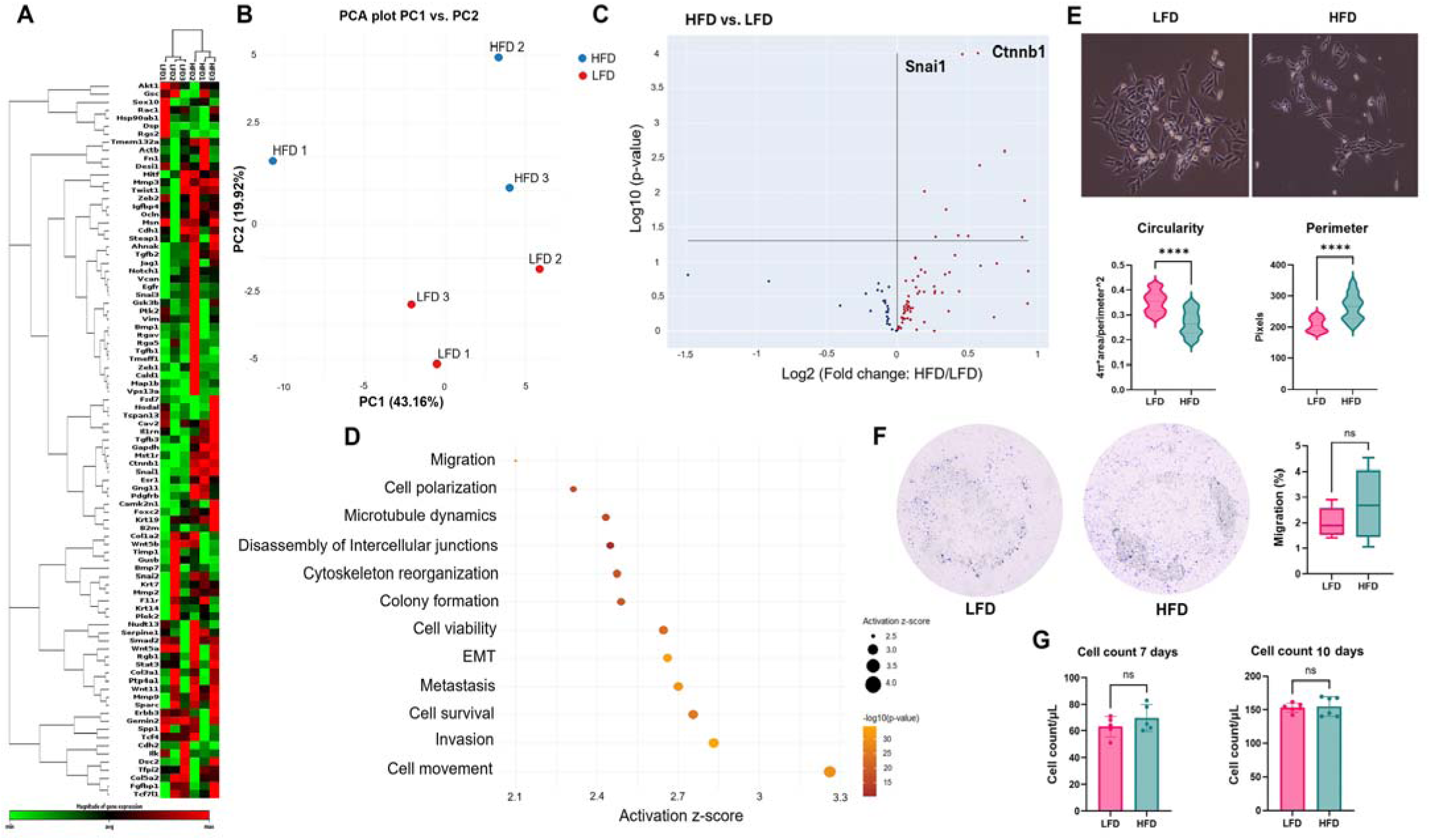
Adipocyte-derived exosomes from HFD-fed mice induce transcription of EMT genes in E0771 cells. E0771 cells were cultured for 3 days with adipocyte-derived exosomes of either HFD or LFD fed mice. Expression of selected EMT genes was analyzed by commercial PCR array and Ingenuity Pathway Analysis (IPA). (A) Heatmap of EMT genes showing differential expression. (B) Principal Component Analysis (PCA) showing clustering of triplicates of the same group regarding EMT gene expression. (C) Volcano plot of significantly upregulated genes (*e.g*., *Snai1*, *Ctnnb1*) in HFD treated group versus LFD. (D) IPA enrichment analysis represented as bubble plot and horizontal bar chart for different cell functions, where activation score (z-score) above two was considered significant (p-value=0.05) and –log 10(p-value) above 1.3 was considered significant (p-value=0.05). N=3. (E) Morphological analysis for cellular perimeter and circularity were measured in each group; one representative image is shown, out of 30 images collected for each experimental condition. N=3. For analysis, 100 cells randomly selected from each replicate for each group were measured using ImageJ. Data were analyzed by unpaired, two-tailed t-test, with statistical significance presented as: ****p < 0.0001. All data are shown as mean ± SD. (F) Migration of E0771 cells was measured in a 6-hour transwell assay. Cells that reached the distal side of the 8-μm pore membrane were visualized by crystal violet and microscopy. Quantitative data are from five independent experiments (N=6). (G) Cell counts were performed at 7 and 10 days by hemocytometer and microscopy (N=6). Data were analyzed by unpaired, two-tailed t-test, with statistical significance presented as: ns, not significant. All data are shown as mean ± SD.

### Plasma-Derived Exosomes Induce EMT and Enhance Cell Migration and Survival

Further investigation into exosomes derived from plasma revealed significant differential expression of EMT genes (Supplementary Figure S3A, S3B). Specifically, pro-EMT genes were upregulated while anti-EMT genes were downregulated in cells treated with exosomes from the HFD group (Supplementary Figure S3C). Notably, the pro-EMT gene *Snai2* showed marked upregulation, and the epithelial marker *Krt14* was significantly downregulated (Figure 3A-C). These changes suggest a shift towards a mesenchymal phenotype. Enrichment analysis of the differentially expressed genes indicated a negative enrichment for anoikis and cell death pathways, suggesting enhanced cell survival. Conversely, pathways related to cell movement, cytoskeleton organization, and cell viability were the most significantly enriched (Figure 3D). This enrichment supports the observation that HFD plasma-derived exosomes promote cellular changes conducive to migration and survival, and these pathways are related to activation of Rac1 and STAT3 networks (Supplementary Figure S3D). Morphological analysis demonstrated that HFD-treated cells exhibited a more mesenchymal-like morphology compared to controls, characterized by elongated cell shapes and increased cytoskeletal reorganization, indicative of a pro-migratory phenotype (Figure 3E). This morphological shift was further corroborated by migration assays, which confirmed increased cell motility in HFD-treated cells (Figure 3F). Additionally, cell viability assays revealed higher cell counts in HFD-treated cells after 7 and 10 days of a single exosome treatment (Figure 3G). This increased viability suggests that exosomes from HFD mice not only promote migration but also enhance cell survival or proliferation. Given these results showing functional differences compared to the adipocyte-derived exosomes, we decided to proceed with plasma-derived exosomes for the rest of the experiments, since they elicit stronger signals in our model.

**Figure 3.**
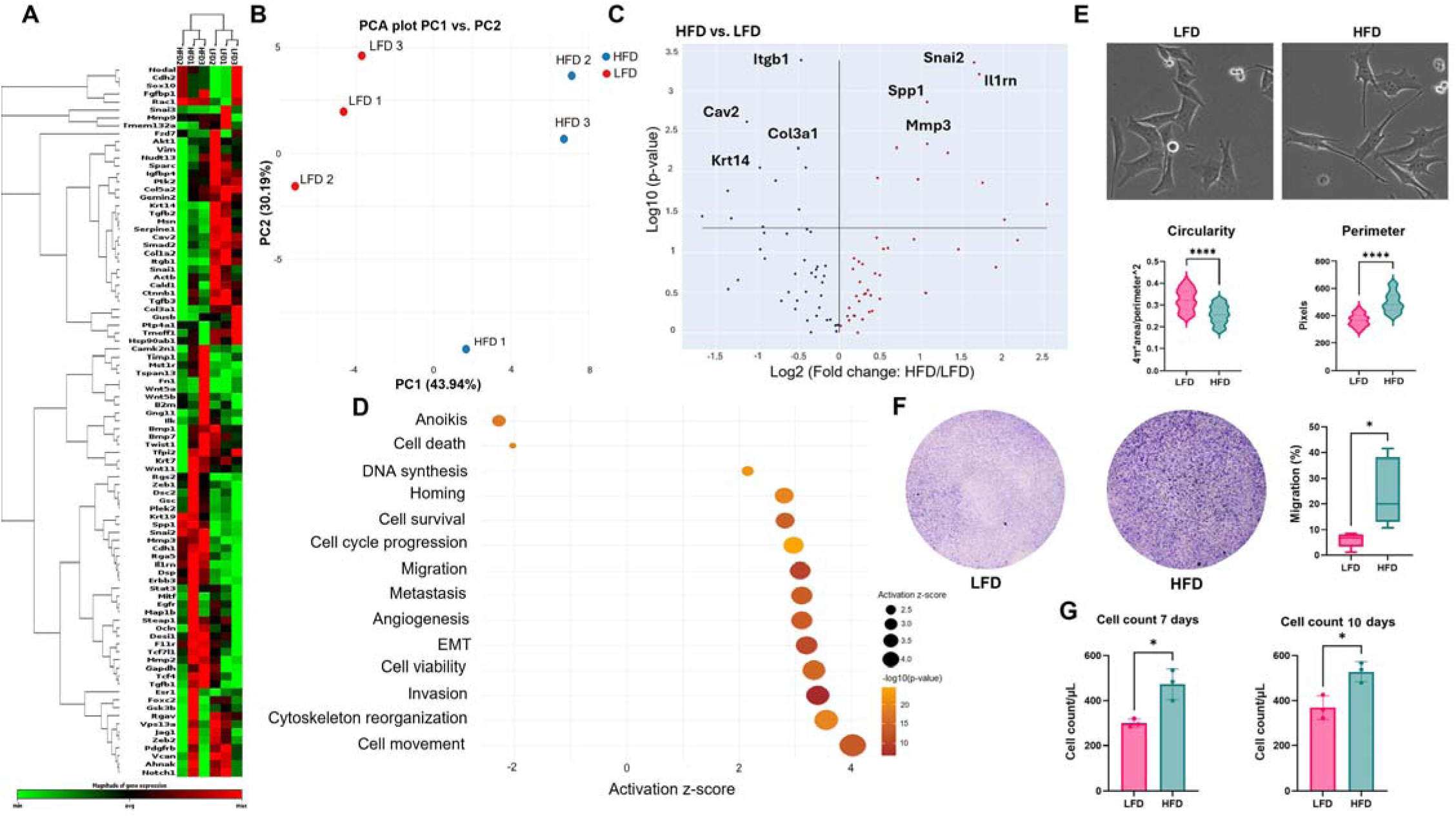
Plasma-derived exosomes from HFD-fed mice induce transcription of EMT genes in E0771 cells. E0771 cells were cultured for 3 days with plasma-derived exosomes of either HFD or LFD fed mice. Expression of selected EMT genes was analyzed by commercial PCR array and IPA. (A) Heatmap of EMT related genes showing differential expression. (B) PCA showing clustering of triplicates of the same group for EMT gene expression. (C) Volcano plot showing significantly upregulated genes (*e.g., Snai2*, *Mmp3*) and downregulated genes (*e.g., Itgb1*, *Krt14*) in HFD treated group versus LFD. (D) IPA enrichment analysis represented as a bubble plot and horizontal bar chart for different cell functions, where activation score (z-score) above two was considered significant (p-value=0.05) and –log 10(p-value) above 1.3 was considered significant (p-value=0.05). N=3. (E) Morphological analysis for cellular perimeter and circularity were measured in each group; one representative image is shown, out of 30 images collected for each experimental condition. N=3. For analysis, 100 cells from each replicate for each group were randomly measured using ImageJ. Data were analyzed by unpaired, two-tailed t-test, with statistical significance presented as: \*\*\*\**p* < 0.0001. All data are shown as mean ± SD. (F) Migration of E0771 cells was measured in a 6-hour transwell assay. Cells that reached the distal side of the 8-μm pore membrane were visualized by crystal violet and microscopy. For quantitation, data are from six independent experiments (N=6). (G) Cell counts were performed at 7 and 10 days by hemocytometer and microscopy (N=3). Data were analyzed by unpaired t-test, two-tailed t-test, with statistical significance presented as: \**p* < 0.05. ns, not significant. All data are shown as mean ± SD.

### Proteomic and Phosphoproteomic Profiling of E0771 Cells Treated *in vitro* with Plasma-Derived Exosomes

After observing a stronger effect on cell migration from plasma-derived exosomes, compared to adipocyte-derived exosomes, we proceeded to analyze the proteomic profile of the exosomes to elucidate potential mechanisms underlying the increased cell migration. Proteomic and phosphoproteomic analyses were conducted on samples from HFD and LFD groups to identify differentially expressed proteins and phosphorylation events (Figure 4A, 4B). The analysis revealed distinct expression profiles between the HFD and LFD groups, with notable differences in proteins involved in cell migration and signaling pathways. The complete dataset of pathway enrichment predicted by gene ontology biological process analysis, comparing HFD to LFD conditions, is shown in Supplementary Table T2. The full dataset of peptide and phosphopeptide identifications is shown in Supplementary Table T3. This analysis highlighted significant enrichment in cell migration pathways, particularly those dependent on the Rho-GTPases signaling cascade (Figure 4C, 4D). Among these, Rac1 was identified as one of the most enriched activation pathways. Rac1 is a well-known regulator of cell motility, cytoskeletal dynamics, and cellular migration, suggesting that its activation may be a key driver of the observed migratory behavior in cells treated with plasma-derived exosomes from the HFD group. Further investigation into the phosphoproteomic data revealed increased phosphorylation of Rac1 and other components of the Rho-GTPases pathway, indicating an upregulation of this signaling cascade in the HFD group. Phosphoproteomic analysis also revealed significant post-translational activation of EMT-related regulators in HFD-exosome-treated cells. Among the phosphorylated residues detected, vimentin showed a significant phosphorylation at S39, this phosphorylation event promotes in vivo tumor metastasis^22^ and increases the ability of vimentin to induce motility and invasion^23^. ZEB1 was also phosphorylated at S682, suggesting enhanced transcriptional repression of epithelial markers^24^. Lastly, Paxillin phosphorylation at S83 was detected, a site previously linked to Rac1 activation and cytoskeletal remodeling via ERK signaling, processes already observed in our experiments and that are important for cancer metastases^25^ (Supplementary Figure S4A). To confirm total Rac protein levels, we performed immunoblotting analysis, showing no differences between LFD and HFD treated E0771 cells (Supplementary Figure S4B), and directly assessed Rac1 pathway activation with a pull-down assay which revealed numerically increased Rac1-GTP levels in cells treated with HFD-derived exosomes compared to LFD controls (Supplementary Figure S4C) ultimately confirming that the observed differences were due to increased activation rather than total protein expression. Consistent with Rac1 role in cytoskeletal remodeling, phalloidin staining of E0771 cells revealed enhanced actin stress fiber formation in the HFD-treated group, including increased total and average branch length, number of branches, and number of junctions per cell (Supplementary Figure S5A-B). These findings suggest that plasma-derived exosomes from obese mice may promote cell migration through the activation of Rac1 and related pathways.

**Figure 4.**
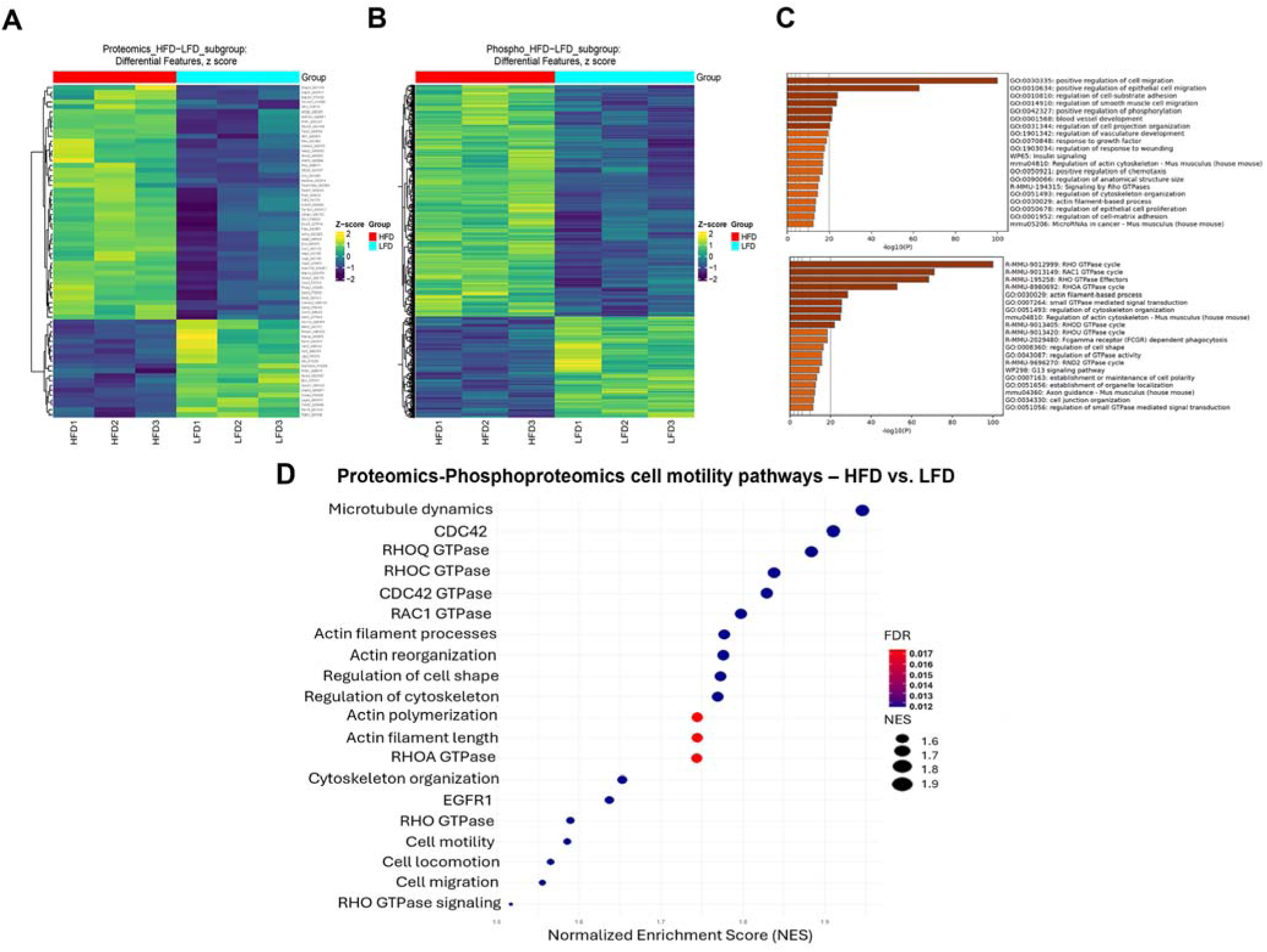
Proteomics and Phosphoproteomics analysis of E0771 cells treated with plasma-derived exosomes. (A) Proteomics and (B) phosphoproteomics heatmaps showing differential expression features between HFD and LFD groups. (C and D) Enrichment analysis identified biological pathways significantly impacted by the differentially expressed proteins. Bubble plot and horizontal bar chart with NES and –log 10(p-value) above 1.3 considered significant (p-value=0.05). N=3.

### Role of Rac1 in the Mesenchymal and Migratory Phenotype Induced by HFD Exosomes

Given the *in vitro* effects of HFD exosomes on E0771 cells and the proteomic analysis suggesting a potential mechanism, we investigated the role of Rac1 in the observed mesenchymal phenotype. Cells treated with either HFD or LFD plasma-derived exosomes for 3 days and subsequently with the Rac1 inhibitor, EHT 1864, were tested for the migratory capabilities in a wound-healing assay. Cells were subjected to a dose-response curve for 24 hours. Where EHT 1864 was used in a ranging concentration between 5-50 µM (Supplementary Figures S6A-C). E0771 cells treated with HFD plasma-derived exosomes and EHT1864 showed an increase in migration and wound closure compared to the LFD group (Figure 5A). A dose-response curve was plotted to determine the IC50 values for both groups, where the IC50 for the LFD group is 26.34 µM and HFD group above 50 µM, observing an increase as well in the area under the curve for the wound closure of the HFD group (Figure 5B). These differences became more evident the higher the concentration of the Rac1 inhibitor was, as shown in the representative images for each group (Figure 5C). We further validated the dependence of HFD-treated E0771 cells to Rho pathways, inhibiting Rac1 and ROCK with EHT-1864 and Y-27632 respectively. We performed cell count and cell shape analysis upon Rac1 and ROCK inhibition. Quantitative imaging revealed increased circularity and decreased perimeter, indicative of reduced cytoskeletal polarization, with significant effects on cell numbers, major differences in cell shape where observed upon Rac1 inhibition rather than ROCKi (Supplementary Figure S6D). Overall, these results confirm that Rho pathways, specifically Rac1, activity is essential for maintaining the migratory morphology induced by HFD-derived exosomes and that its inhibition effectively reverts this phenotype, potentially contributing to the aggressive phenotype observed in cancer cells.

**Figure 5.**
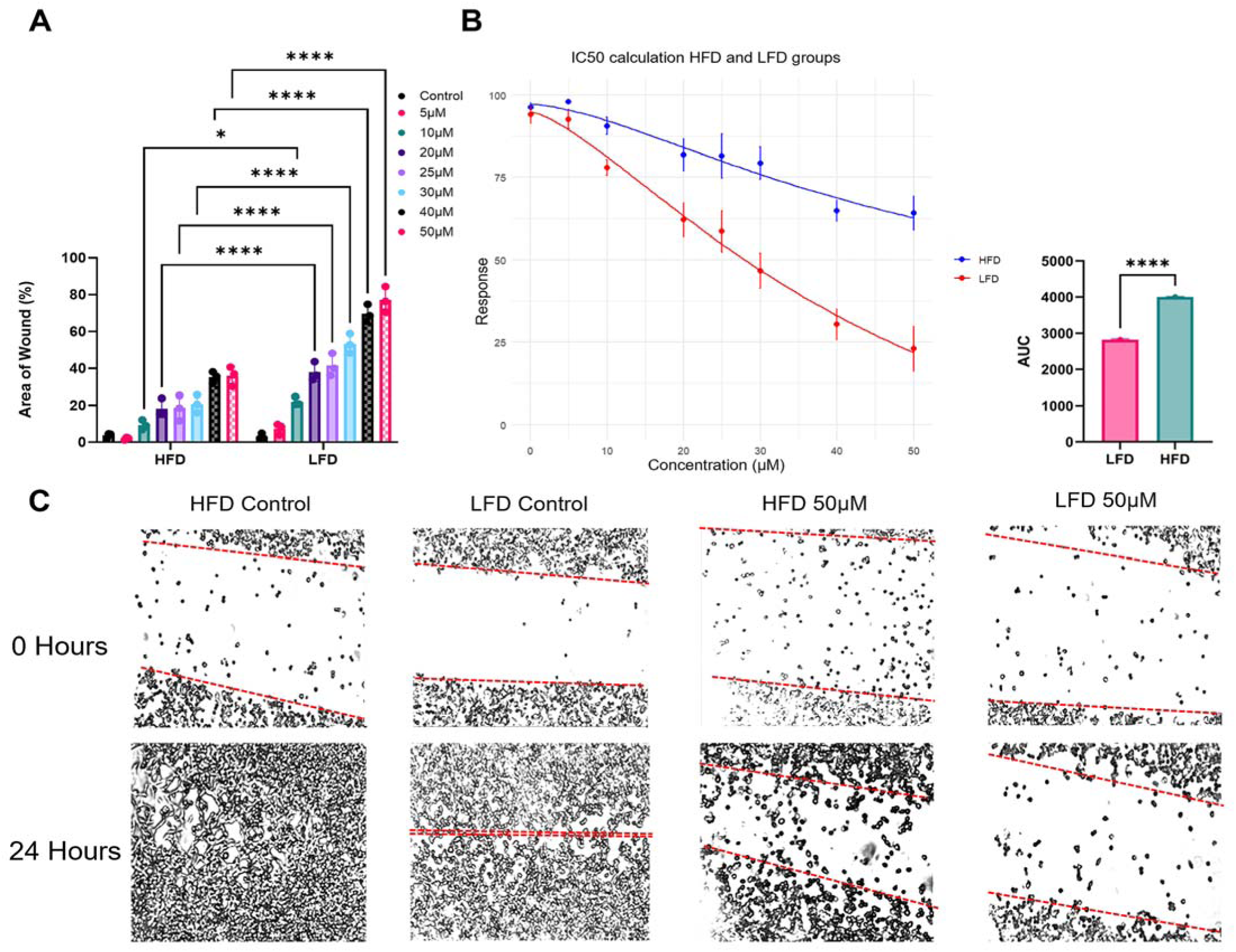
Rac1 role in E0771 cells treated with plasma-derived exosomes. E0771 cells were cultured for 3 days with plasma-derived exosomes of either HFD or LFD fed mice, after these 3 days, cells were treated with an increasing concentration of the Rac1 inhibitor, EHT1864. Concentrations ranged 5-50 µM, leaving one group as a control without Rac1 inhibitor. (A) Area of the wound represented as a percentage of the total area in time=0. The area of the wound left after 24 hours of the scratch is lower for the cells that were treated with plasma-derived HFD exosomes, indicating and increase in the closure of the wound. Data were analyzed by 2-way Anova with a Šidák’s post-hoc analysis for multiple comparison testing with statistical significance presented as: *p < 0.05; ****p < 0.0001. All data are shown as mean ± SD. (B) Dose-response curve for different EHT1864 concentrations (0, 5, 10, 20, 25, 30, 40 and 50 µM). The area under the curve between both groups shows a statistical difference between the level of response to EHT1864 inhibitor. Data were analyzed by unpaired, two-tailed t-test, with statistical significance presented as: ****p < 0.0001. All data are shown as mean ± SD. (C) Representative images of the wound-closure assay at 0 hours and 24 hours after the scratch. These images show the differential closure between HFD and LFD at high concentrations of EHT 1864. N=3.

### Enhanced Metastatic Potential of E0771-GFP Cells Treated with HFD Plasma-Derived Exosomes

To investigate the aggressive mesenchymal phenotype in an animal model, E0771-GFP cells treated with LFD or HFD plasma derived-exosomes for three days were injected into the left second mammary fat pad of six-week-old female C57BL/6J mice. After 28 days, the mice were sacrificed, and the brain and lung tissues were analyzed (Figure 6A). Both the lung and brain tissues were homogenized, seeded in cell culture plates, and allowed to grow for 7 days (Figure 6B). Subsequently, we assessed the number of E0771-GFP colonies (colony count), their size (colony coverage), and their cell density (cell count). Our analyses revealed that both the brain and lung tissues showed a higher number and larger size of colonies in the HFD group compared to the LFD group. Specifically, the brain tissue exhibited a higher cell density in the HFD group, while the lung tissue showed no significant differences in cell density between the groups Notably, in the LFD group, only two out of five mice had brain metastases, whereas all mice in the HFD group showed the presence of E0771 cells in the brain (Supplementary Table T4). However, mice pretreated with HFD exosomes exhibited a significantly higher burden of brain and lung metastases compared to the LFD group (Figure 6C). These findings suggest that exosomes derived from HFD conditions enhance the metastatic potential of E0771-GFP cells, particularly promoting brain and lung metastasis.

**Figure 6.**
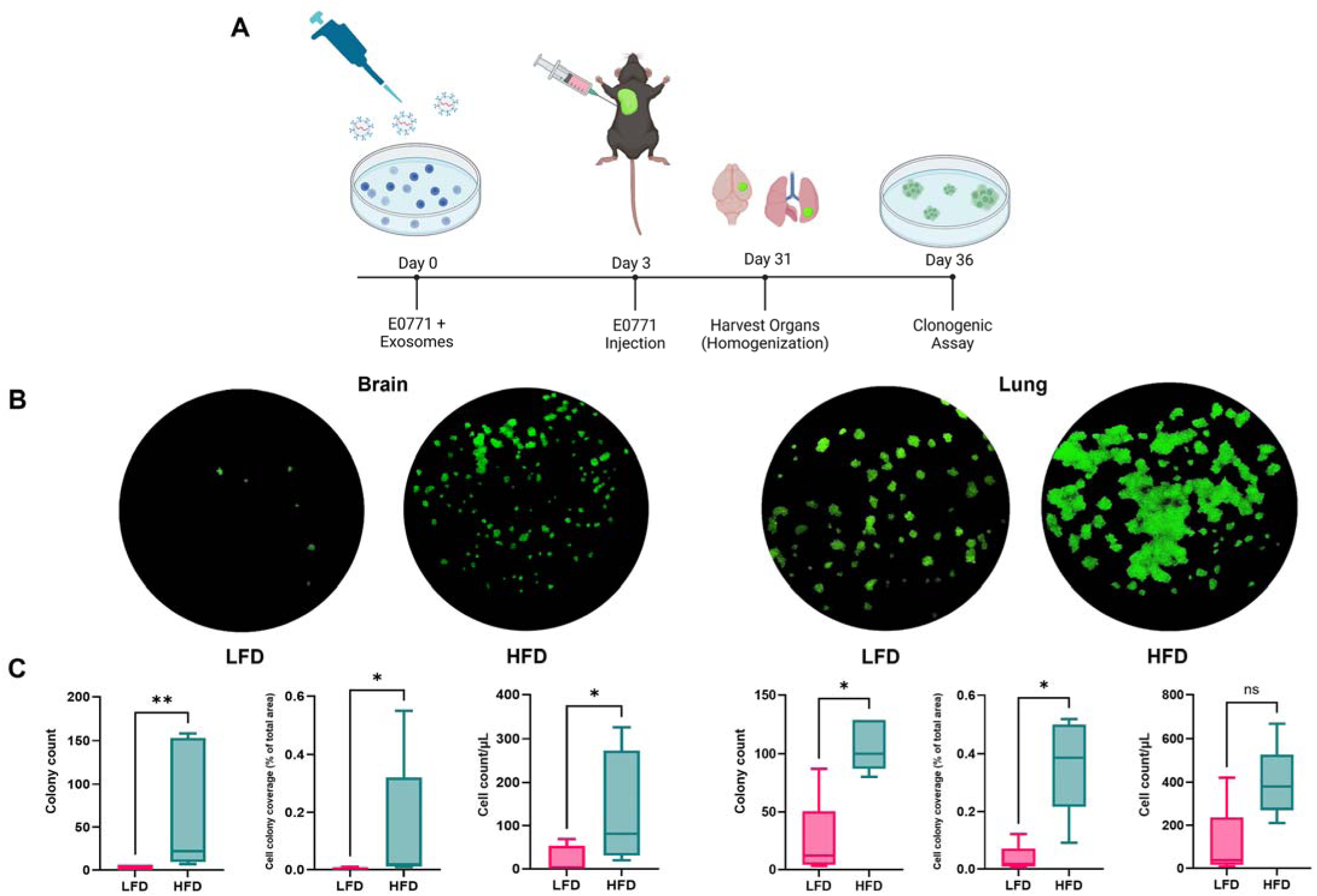
Plasma-derived exosomes from HFD-fed mice promote metastasis burden in brain and lung of C57BL/6J female mice. E0771 cells were cultured for 3 days with plasma-derived exosomes of either HFD or LFD fed mice, then 100,000 of these cells were injected in the 2nd mammary fat pad of 6-week-old C57BL/6J female mice. Metastases burden was measure by clonogenic assay. (A) Schematic representation of the hybrid in vitro and in vivo experiment. Image generated with BioRender® software (BioRender.com). (B) Fluorescent microscopy of a chosen representative 10cm2 tissue culture plate of a clonogenic assay for both HFD and LFD group and for brain and lung homogenized and tissues (N=5). (C) Statistical comparison for three different parameters accounting for metastasis burden. E0771-GFP cells treated with HFD plasma-derived exosomes show a significant increased metastasis burden compared to the control group (LFD). Data were analyzed by Mann-Whitney test with statistical significance presented as: ns, not significant, *p < 0.05; **p < 0.01. All data are shown as mean ± SD. N=5.

### Transcriptome Analysis of *in vitro* E0771 Cells and Brain Metastatic Cells

Given the significant differences in the brain metastasis burden between HFD and LFD exosome-treated cells. We decided to run Bulk RNA-sequencing on *in vitro* cells 3 days after treatment and before injection in the 2^nd^ mammary fat pad, plus the cells that were obtained from brain metastasis after the 28-day period after injection in the mice. We found a great variety of differentially expressed genes in both brain mets (Figure 7A) and cells *in vitro* (Figure 7B) when comparing HFD to LFD treated E0771 cells. Due to the lower incidence of brain metastasis in the LFD treated group, we included a negative control group where E0771-GFP cells were not treated with exosomes to reach an N=5. As expected, LFD (represented as LB group) treated cells and non-exosome treated cells (represented as CB group) clustered together for brain metastatic cells while HFD treated group (represented as HB group) clustered together by itself. Since we observed the LFD treated group was like the negative control, we decided to run enrichment analysis to have a higher power analysis and cluster LFD plus negative control as a one control group versus the HB group. Hallmark gene set enrichment analysis was performed using RNA-seq data from cells treated *in vitro* and from the ones injected into the mammary fat pad of mice that later metastasized to the brain (Figure 7C). The hypergeometric test revealed significant enrichment of several hallmark gene signatures in both the brain metastasis and *in vitro* conditions, particularly after HFD plasma-derived exosome treatment, as compared to the control groups. *In vitro*, the early estrogen response was significantly downregulated, while in brain metastases, the late estrogen response showed marked downregulation. Additionally, both the glycolysis and hypoxia pathways were notably upregulated in the brain metastasis group following HFD treatment. Immune-related pathways also exhibited significant changes, with both the inflammatory response and interferon alpha/gamma signaling showing strong enrichment in the brain metastasis samples. Finally, mTORC1 signaling was upregulated in brain metastases under HFD conditions, while TNF-alpha signaling via NF-kB was enriched in the *in vitro* cells. To further explore the clinical relevance of these findings, we performed survival analysis using data from TCGA (Figure 7D) breast cancer cohorts. Expression levels of genes identified in our study were categorized into high and low expression groups, adjusted for clinical variables such as age and proliferation rates.Kaplan-Meier survival curves from the TCGA cohort demonstrated a significant difference in overall survival between patients with high and low expression of these genes (p = 0.006), with high expression associated with worse survival outcomes. These findings suggest that the transcriptional consequences of exosomal programming in our mouse model are clinically relevant to aggressive TNBC in humans, and highlight the potential clinical impact of metabolic and immune pathways in brain metastasis, particularly in the context of HFD plasma-derived exosome treatment.

**Figure 7.**
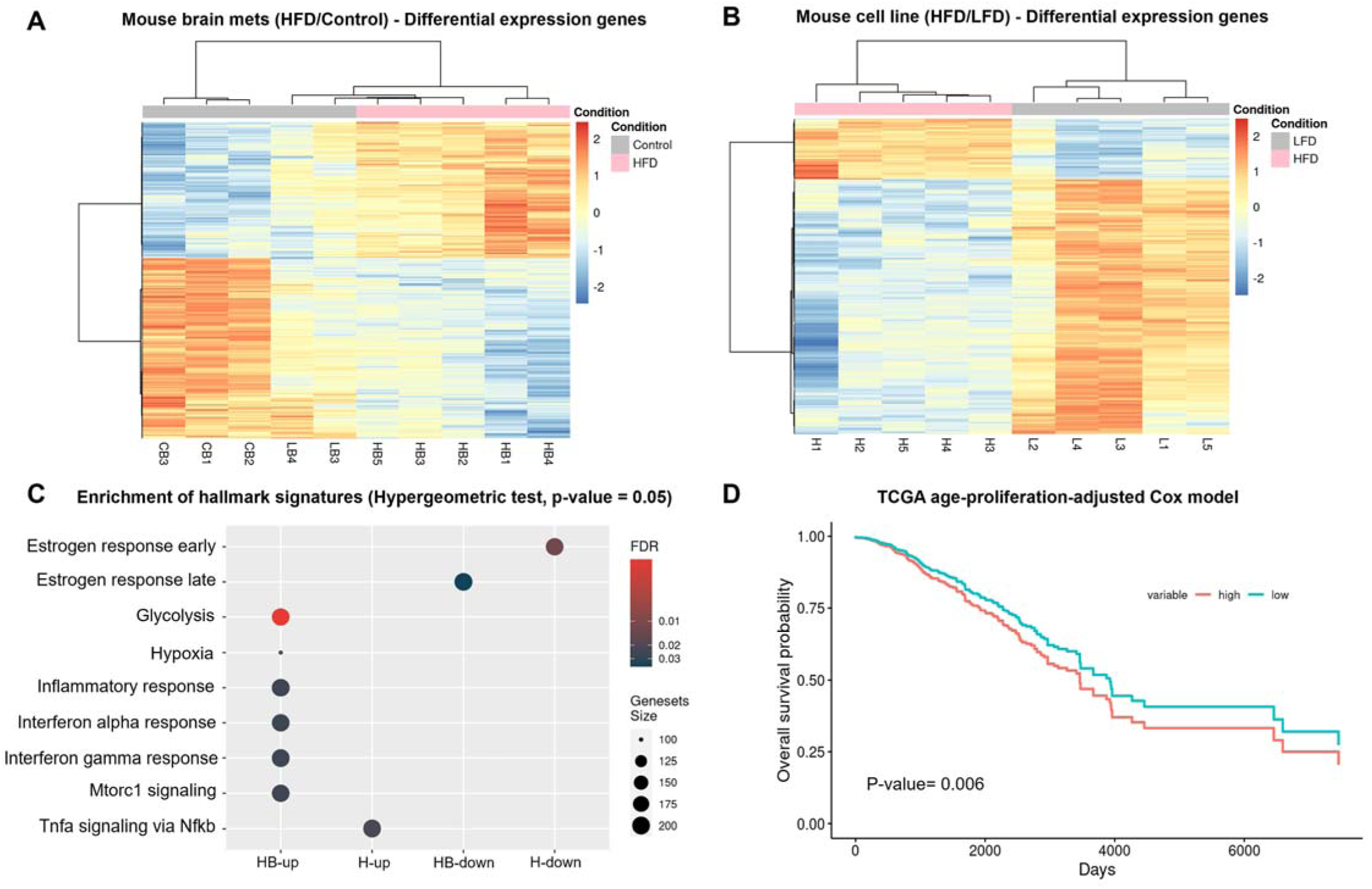
Transcriptome analysis of *in vitro* and brain metastatic E0771 cells. **(A)** Heatmap showing differential gene expression in brain metastatic cells between HFD and control groups. **(B)** Heatmap of differential expression features in E0771 cells cultured *in vitro* after HFD and control treatments. (C) Enrichment of hallmark gene signatures showing pathways significantly impacted by differential mRNA expression following treatment. (D) TCGA BRCA-specific survival analysis. Age and Proliferation Adjusted Survival curve of mouse brain metastases (HFD/LFD) showing overall survival probability, with high and low gene expression groups. The difference between groups is statistically significant (P-value = 0.006).

## DISCUSSION

Population studies over the past twenty years have strongly implicated obesity and diabetes as risk factors in the incidence and progression of obesity-driven cancers, including breast cancer^26,27^. Although much of the early epidemiological literature linking obesity and breast cancer has focused on ER-positive tumors, more recent studies have begun to explore the impact of metabolic dysfunction in ER-negative subtypes such as TNBC. Obesity and T2D are increasingly recognized as contributors to poorer breast cancer outcomes across subtypes. In a landmark prospective study, Calle et al. reported that obesity accounts for approximately 20% of all cancer-related deaths in women in the U.S., including breast cancer^28^.Renehan et al. extended these findings in a large meta-analysis, associating higher BMI with increased incidence of breast cancer, though primarily among postmenopausal women ^29^. Importantly, studies such as the Black Women’s Health Study have begun to stratify breast cancer mortality by both the estrogen receptor status and duration of T2D. They observed a notably higher risk of mortality in women with long-standing T2D, suggesting that metabolic dysfunction may facilitate systemic disease progression independent of hormone receptor expression^30^. Recent studies have shown that obesity-induced metabolic dysfunction and adipocyte senescence can alter exosomal signaling in ways that promote tumor progression. Li et al. demonstrated that obesity and hyperinsulinemia drive adipocyte senescence through activation of cell cycle programs^31^, while Ishaq et al. showed that caloric restriction reverses senescence and inflammation related features in adipose tissue, supporting the idea that metabolic interventions may restore exosomal homeostasis in obesity-driven disease contexts^32^. These findings are consistent with our own data showing that HFD-derived exosomes enhance migratory and mesenchymal features in TNBC cells and promote metastasis to distant organs, including the brain. Collectively, this literature reinforces the mechanistic relevance of our obesity-exosome-TNBC model and supports the broader hypothesis that extracellular vesicles act as key metabolic messengers in cancer progression.

These findings underscore the urgent need to elucidate how systemic metabolic alterations mechanistically influence TNBC progression. Our study addresses this gap by identifying exosomes from insulin-resistant, obese mice as functional mediators capable of promoting EMT and enhancing metastatic traits in TNBC cells. This mechanistic insight helps contextualize the epidemiologic associations and supports the notion that targeting metabolic-exosomal signaling could yield novel therapeutic strategies for aggressive breast cancer in metabolically vulnerable populations.

However, the key causal elements that connect obesity-driven, systemic insulin resistance to increased metastatic burden and tumor progression remain poorly understood. As other groups have described^33,34^, the growth of E0771 tumors is enhanced in obese mice compared to lean mice, and we have separately shown that plasma or adipocyte derived exosomes that reflect diabetic or insulin resistant features of obesity strongly promote tumor aggressiveness in multiple cancer types^14,15,16^ which led us to hypothesize that exosomes either present in plasma or derived from adipocyte exosomes promote tumor growth and aggressiveness. In this study, we utilized an HFD-fed C57BL/6J mouse model to investigate the impact of obesity-driven insulin resistance on exosome production and function, focusing particularly on the role of exosomes in promoting a mesenchymal phenotype in E0771 cells as a model for triple negative breast cancer in women with obesity-driven metabolic disease.

Our results demonstrate that HFD-induced obesity and insulin resistance significantly alter exosomal content, leading to profound molecular and phenotypic changes upon uptake into breast cancer cells, compared to lean and insulin sensitive controls. We first demonstrated that the HFD-induced obesity model was successfully established, evidenced by standard methods of validation like oGTT and ipITT to confirm systemic insulin resistance, by showing abnormalities in glucose clearance and insulin responsiveness respectively^35,36^, significant weight gain and increased body weight^37^.

Exosomes isolated from HFD-treated mice promoted an EMT signature of breast cancer cells distinct and significantly worse than LFD controls. EMT array analysis revealed upregulation of key EMT markers, such as Snai1 and Ctnnb1, and morphological changes indicative of mesenchymal phenotype and enrichment of pathways related to cytoskeleton organization, migration, and cell survival. This pattern has been already reported by other groups, where cell morphology and migration are linked to upregulation of EMT genes, including the genes mentioned above^38,39^. Despite these molecular changes, migration assays showed no functional difference in migratory capacity under the tested conditions. This discrepancy suggests that whereas HFD-derived exosomes induce molecular markers of EMT, additional factors or conditions are required to translate these transcriptional changes into enhanced migratory behavior.

Further analysis of plasma-derived exosomes from HFD-treated mice showed significant upregulation of pro-EMT genes and downregulation of epithelial markers, indicating a shift towards a mesenchymal phenotype. These exosomes enhanced cell motility, viability and migratory capability, suggesting a more pronounced role in promoting cancer progression compared to adipocyte-derived exosomes. These findings underscore the importance of the source of exosomes and suggest that plasma-derived exosomes have a more significant role in promoting aggressive cellular behaviors.

Proteomic and phosphoproteomics analyses further highlighted the involvement of Rac1 and the Rho-GTPase signaling cascade, which is a known regulator of cell motility and cytoskeletal dynamics^40,41,42^. The upregulation of Rac1 and Rho-GTPase pathways aligns with increased cell motility and mesenchymal transition, suggesting that these signaling pathways are potential mediators of the pro-EMT effects induced by HFD exosomes. This was further supported by Rac1 pull-down assays, which showed higher numerical levels of active Rac1 in HFD-treated cells relative to LFD and HFD + EHT1864 groups, confirming the trend observed in the phosphoproteomic analysis. Concordantly, phalloidin staining revealed increased actin stress fiber formation and cell shape changes, reinforcing the functional activation of Rac1-dependent cytoskeletal remodeling.

*In vivo* experiments using E0771-GFP cells treated with HFD or LFD plasma-derived exosomes demonstrated a higher metastatic burden in the brain and lungs of mice under the HFD condition. These results are consistent with previous clinical studies, where patients with TNBC and comorbid obesity face increased risk of developing secondary tumors in the brain^43^, suggesting that plasma exosomes may enhance the brain metastatic potential of breast cancer patients with obesity. The greater number and size of colonies in the brain and lungs in our model, particularly the universal presence of brain metastases in the HFD group, highlight the aggressive nature of the cancer cells promoted by these exosomes.

Epidemiological and clinical studies have firmly established obesity and lack of physical activity as modifiable risk factors that significantly influence breast cancer progression. As mentioned before, Calle et al. reported that excess body weight accounts for a substantial proportion of cancer-related deaths in the U.S. while Renehan et al confirmed this positive association between BMI and postmenopausal breast cancer incidence across multiple cohorts^28,29^. On the other hand, exercise has been shown to influence tumor biology through multiple mechanisms, including improving insulin sensitivity, reducing systemic inflammation, and modulating the tumor microenvironment via effects on immune function, oxidative stress, and epigenetic regulation^44^. Recent work has focused on extracellular vesicles as direct mediators of exercise’s anti-tumorigenic effects. Mlynska et al, demonstrated that extracellular vesicles isolated from exercised mice attenuate TNBC progression and promote CD8+ T cell recruitment^45^. Similarly, Puurand et al. demonstrated that exercise-induced extracellular vesicles carry bioactive cargo, including microRNAs and proteins, which can modulate cancer cell metabolism by suppressing glycolysis and enhancing oxidative phosphorylation, potentially reducing metastatic potential^46^. Our data position obesity-derived exosomes as active agents of cancer progression and set the stage for future experiments assessing whether exercise-derived exosomes may counteract these pro-metastatic signals.

RNA-seq results highlight downregulation of early estrogen response *in vitro* and late estrogen response in brain metastases, consistent with the ER-negative nature of E0771 cells^47^. This pattern suggests the cells have become estrogen-independent, characteristic of TNBC. Additionally, significant enrichment of glycolysis and hypoxia pathways in the brain metastases points to metabolic reprogramming^48,49^, where tumor cells adopt aerobic glycolysis to survive in the nutrient-deprived and hypoxic brain environment, a key adaptation that might favor E0771 cells treated with HFD plasma-derived exosomes to colonize and create a higher metastatic burden in brain compared to the control group. Survival analysis using the TCGA breast cancer cohort further underscores the clinical relevance of these metabolic and immune alterations, with high expression of the genes identified in the mouse model being significantly associated with worse overall survival outcomes among patients. This finding suggests that the pathways modulated by HFD-derived exosome treatment in our mouse models have translational relevance to certain subtypes of human breast cancer, particularly those captured in TCGA. It is important to note that, while TNBC is not explicitly labeled in TCGA, we focused our analysis on the basal-like molecular subtype, which is widely accepted as a close transcriptomic proxy for TNBC. Our HFD-induced metastasis gene signature showed a significant negative association with overall survival in this group, reinforcing the translational applicability of our preclinical model. These findings suggest that obesity-associated exosomal programming may drive molecular phenotypes linked to poor prognosis in basal-like breast cancer, potentially through modulation of metabolic and immune-related pathways.

Overall, our RNA-seq analysis identified metabolic and immune-related changes, such as enrichment of inflammatory response and interferon alpha/gamma signaling, between HFD and LFD exosome-treated cells. These findings should be validated in more complex *in vivo* systems. Given this focus on immune modulation, the pathway directionality was not strong nor specific.

However, proteomics analysis revealed robust activation of Rac1 and Rho GTPase signaling, which are crucial for cytoskeletal reorganization, migration and metastasis. In addition, Rho GTPases are thought to play a role in establishing an immunosuppressive tumor microenvironment^50^, which links both observations. In the setting of obesity and insulin resistance, Rho GTPases contribute to glucose homeostasis by performing critical metabolic functions within skeletal muscle and adipose tissue^51,52^. Together with the EMT gene array data and enhanced mesenchymal morphology, these phospho-modifications reinforce a robust activation of EMT through both transcriptional and post-translational mechanisms. These findings led us to prioritize these pathways for further investigation, given their established role in driving tumor invasion and brain metastasis. Further investigation into the role of Rac1 activation revealed its critical involvement in promoting the mesenchymal and migratory phenotype observed in HFD-treated cells. The use of the Rac1 inhibitor EHT-1864^53,54^ demonstrated dose-dependent inhibition of migration and wound closure, with HFD-treated cells exhibiting higher resistance to Rac1 inhibition.

The role of Rho GTPases, particularly Rac1, in cancer progression is increasingly recognized as a critical factor in driving breast cancer metastasis^55^. Moreover, ongoing clinical trials, currently in Phase 1, investigate the use of oral MBQ-167 (a compound targeting Rho GTPases) as a single agent for participants with advanced breast cancer^56^. Therefore, Rho GTPases and Rac1 could serve as critical mediators linking the metabolic dysfunction caused by obesity and systemic insulin resistance with the increased metastatic potential of TNBC. This understanding positions Rac1 as a promising therapeutic target, particularly in patients with obesity-related metabolic comorbidities, providing a rationale for further exploration of Rac1 inhibition in mitigating metastatic burden. Considering the large burden of obesity and metabolic disease among adult Americans, with >120 million people currently diagnosed as diabetic or pre-diabetic^57^, the public health significance of these insights for breast cancer patients is apparent.

Numerous studies now support the role of obesity-derived exosomes in driving aggressive breast cancer phenotypes, particularly in the context of TNBC. Exosomes isolated from obese adipose tissue or plasma have been shown to carry distinct cargo, such as oncogenic miRNAs and pro-inflammatory cytokines that modulate tumor behavior through both local and systemic^58,59^. Barone, et al demonstrated that miRNAs contained in exosomes from obese adipocytes enhance epithelial-to-mesenchymal transition and tumor invasiveness in TNBC models^60^. In our group, we have also identified obesity-associated exosomal miRNAs, such as miR-155 and miR-27a, as key regulators of cell migration, glycolysis, and immunosuppression within the tumor microenvironment^14^.

Our supplementary analysis reveals distinct miRNA profiles in plasma-derived exosomes from HFD and LFD mice, underscoring the impact of metabolic status on exosomal content (Supplementary Figure S7A). HFD-derived exosomes show increased levels of miR-192-5p, miR-12182-3p, miR-122-5p, and miR-122b-3p, whereas LFD-derived exosomes are enriched in miR-184-3p, miR-125a-5p, and miR-485-5p. As previously described, our group has identified miRNAs as the main effectors driving the observed cellular responses to exosomal treatment^15,16^. This differential miRNA expression activates distinct pathways and mechanisms, as illustrated in the HFD *vs*. LFD GO enrichment plot (Supplementary Figure S7B). The enrichment bubble plot emphasizes high gene ratios in pathways associated with GTPase binding, cell leading edge formation, filopodium structure, and actin-based cell projections, among others. These findings suggest that the miRNAs enriched in HFD exosomes contribute to cytoskeletal dynamics and cell survival mechanisms, which are crucial for enhanced motility and invasive potential. This interpretation aligns closely with other data presented in this study, specifically, the observed changes in cell migration, morphology, and involvement of Rho GTPases.

Our study provides compelling evidence that exosomes derived from HFD-fed mice significantly promote breast cancer progression and metastasis. These findings underscore the critical role of metabolic conditions in modulating exosomal content and their subsequent impact on cancer cell behavior. The differential effects observed between exosomes derived from visceral adipose tissue and plasma suggest that circulating exosomes may have a more pronounced impact on promoting a mesenchymal and metastatic phenotype in TNBC cells. Future research should explore therapeutic strategies targeting specific signaling pathways, such as Rac1, to mitigate the pro-metastatic effects of obesity-driven insulin resistance.

## Abbreviations

ANOVA: analysis of variance
AUC: area under the curve
BMI: body mass index
CSC: cancer stem-like cell
DMEM: Dulbecco’s modified Eagle medium
EDTA: ethylene diamine tetraacetic acid
EMT: epithelial-to-mesenchymal transition
ER: estrogen receptor
FBS: fetal bovine serum
GFP: green fluorescent protein
HER2: human epidermal growth factor receptor 2
HFD: high fat diet
HPLC: high performance liquid chromatography
IACUC: institutional animal care and use committee
IL: interleukin
IR: insulin resistant
ipITT: intraperitoneal insulin tolerance test
IPA: ingenuity pathway analysis
LFD: low fat diet
oGTT: oral glucose tolerance test
PCA: principal components analysis
PCR: polymerase chain reaction
PBS: phosphate buffered saline
RPM: revolutions per minute
T2D: Type 2 diabetes
TCGA: The Cancer Genome Atlas
TNBC: triple negative breast cancer
TNF: tumor necrosis factor.

## Declarations

### Ethics approval and consent to participate

All animal experiments were performed humanely and in accordance with approved procedures, with oversight of the Institutional Animal Care and Use Committee (IACUC) of Boston University Medical Center. Consent to Participate, and Consent to Publish declarations are not applicable.

### Consent for publication

No humans participated as research subjects in this study.

### Availability of data and materials

The datasets generated and analyzed during the current study are publicly available in the following repositories:

- **RNA-sequencing data:** Deposited in the GEO repository under accession number GSE282303. Reviewer token: svclmaoujbwzted
- **Small RNA-sequencing data:** Available in the GEO repository under accession number GSE282305. Reviewer token: sfwpwoggvvshdun
- **Proteomics and phosphoproteomics data:** Accessible in the ProteomeXchange repository via the PRIDE partner repository under accession number PXD057911. Reviewers will be required to log in through the PRIDE portal with a reviewer username. Reviewer access details:
  - Reviewer username:
  - Password: UWLFsHhJZyax

All datasets are fully accessible and can be used for further research as indicated. Reviewers can access the private datasets during the peer review process using the provided links and tokens.

### Authors’ contributions

Pablo Llevenes conceptualized the study, directed the project, performed all experiments, analyzed the data, created the figures, and wrote the manuscript. Andrew Chen processed and analyzed the RNA-seq data from cells treated with exosomes and created the associated plots. Matthew Lawton conducted the proteomics and phosphoproteomics analyses, including sample processing, data analysis, and figure preparation. Alejandro Rondon-Ortiz assisted with the experimental procedures of immunoblots and Rac1 pull-down assay. Yuhan Qiu and Michael Seen provided technical support, experimental assistance, and valuable scientific input throughout the study. Stefano Monti contributed expertise and oversight for RNA-seq data analysis. Gerald V. Denis supervised and conceived the project, provided essential guidance and critical feedback, contributed to the study design and data interpretation, and played a central role in refining the manuscript. All authors reviewed and approved the final manuscript.

## Acknowledgements

We thank Boston University-Boston Medical Center Cancer Center faculty R. Flynn, N. Ganem and Valentina Perissi, for helpful comments and suggestions. We thank Anna Belkina, Shari Brezinsky and Brian Tilton, the BUMC Flow Cytometry Core Facility, Matthew Bo Au of the BUMC Analytical Core Facility; Timothy Padera and Hengbo Zhou for the cell line E0771-GFP; Stanley Goldstein for his assistance in properly formatting and uploading the proteomics data here included to be compliant with repository guidelines; and Nabil Rabhi for his recommendations in animal handling, and procedures including blood extraction, glucose and insulin tests in mice.

## References

1. Giaquinto AN, Sung H, Newman LA, et al. Breast cancer statistics 2024. CA Cancer J Clin. 2024; 1–19.

2. Foulkes WD, Smith, IE, Reis-Filho JS. Triple-negative breast cancer. New England Journal of Medicine 2010; 363(20), 1938–1948.

3. Bianchini G, Balko JM, Mayer IA, Sanders ME, Gianni L. Triple-negative breast cancer: challenges and opportunities of a heterogeneous disease. Nature Reviews Clinical Oncology 2016; 13(11), 674–690.

4. Henry NL, Shah PD, Haider I, Freer PE, Jagsi R, Sabel MS. Chapter 88: Cancer of the Breast. In: Niederhuber JE, Armitage JO, Doroshow JH, Kastan MB, Tepper JE, eds. Abeloff’s Clinical Oncology. 6th ed. Philadelphia, Pa: Elsevier; 2020. Included in: American Cancer Society. (2020). Breast Cancer Survival Rates.

5. Perrone MA, Pierdominici M, Parrinello G, Malorni W, Salerno G. Metabolic diseases and breast cancer: the interplay between tumor microenvironment and cellular metabolism. Cancers 2021; 13(5), 963.

6. Kolb R, Zhang W, DeAngelis, T. Obesity and cancer: inflammation bridges the two. Current Opinion in Pharmacology 2018; 42: 87–102.

7. Park J, Euhus, DM, Scherer PE. Paracrine and endocrine effects of adipose tissue on cancer development and progression. Endocrine Reviews 2011; 32(4): 550–570.

8. Park J, Morley TS, Kim M, Clegg DJ, Scherer PE. Obesity and cancer—mechanisms underlying tumour progression and recurrence. Nature Reviews Endocrinology 2014; 10(8): 455–465.

9. Brown KA, Simpson ER, Rudnicka C. Obesity and breast cancer: mechanisms underlying risk and response to treatment. Frontiers in Endocrinology 2010;1: 26.

10. Sun X, Casbas-Hernandez P, Bigelow C, Makowski L, Joseph Jerry D, Troester MA. Normal breast tissue of obese women is characterized by an inflammatory signature. Breast Cancer Research and Treatment 2012;131(3): 1021–1037.

11. Kalluri R, LeBleu VS. The biology, function, and biomedical applications of exosomes. Science, 2020; 367(6478), eaau6977.

12. Guay C, Kruit JK, Rome S, Menoud V, Mulder NL, Jurdzinski A, … Regazzi R. Lymphocyte-derived exosomes promote β-cell death in type 1 diabetes. Diabetes 2019; 68(9): 1782–1793.

13. Costa-Silva B, Aiello NM, Ocean AJ, Singh S, Zhang H, Thakur BK, … Lyden D. Pancreatic cancer exosomes initiate pre-metastatic niche formation in the liver. Nature Cell Biology 2015; 17(6), 816–826.

14. Jafari N, Kolla M, Meshulam T, Shafran JS, Qiu Y, Casey AN, Pompa IR, Ennis CS, Mazzeo CS, Rabhi N, Farmer SR, Denis GV. Adipocyte-derived exosomes may promote breast cancer progression in type 2 diabetes. Sci Signal. 2021 Nov 23;14(710):eabj2807.

15. Jafari N, Chen A, Kolla M, Pompa IR, Qiu Y, Yu R, Llevenes P, Ennis CS, Mori J, Mahdaviani K, Halpin M, Gignac GA, Heaphy CM, Monti S, Denis GV. Novel plasma exosome biomarkers for prostate cancer progression in co-morbid metabolic disease. Adv Cancer Biol Metastasis. 2022;6:100073.

16. Jafari N, Llevenes P, Denis GV. Exosomes as novel biomarkers in metabolic disease and obesity-related cancers. Nat Rev Endocrinol. 2022;18(6):327–328.

17. Patel H, Ewels P, Manning J, Garcia MU, Peltzer A, Hammarén R, Botvinnik O, Talbot A, Sturm G, nf-core bot, Zepper M, Moreno D, Vemuri P, Binzer-Panchal M, silviamorins, Pantano L, Syme R, Kelly G, Hanssen F, Fellows Yates JA, Espinosa-Carrasco J, Fenouil R, Zappia L, Cheshire C, Miller E, Hoeppner M, Zhou P, Guinchard S, Gabernet G, Mertes C. nf-core/rnaseq: nf-core/rnaseq v3.15.1 - Augmented Aluminium Axolotl (Version 3.15.1) [Software]. Zenodo. 2024.

18. Love MI, Huber W, Anders S. Moderated estimation of fold change and dispersion for RNA-seq data with DESeq2. Genome Biol. 2014;15:550.

19. Therneau TM. A package for survival analysis in R. R package version 3.5–7. 2023.

20. The Cancer Genome Atlas Network. Comprehensive molecular portraits of human breast tumours. Nature 2012;490:61–70.

21. Curtis C, Shah SP, Chin SF, Turashvili G, Rueda OM, Dunning MJ, Speed D, Lynch AG, Samarajiwa S, Yuan Y, Gräf S, Ha G, Haffari G, Bashashati A, Russell R, McKinney S, METABRIC Group, Langerød A, Green A, Provenzano E, Wishart G, Pinder S, Watson P, Markowetz F, Murphy L, Ellis I, Purushotham A, Børresen-Dale AL, Brenton JD, Tavaré S, Caldas C, Aparicio S. The genomic and transcriptomic architecture of 2,000 breast tumours reveals novel subgroups. Nature 2012;486:346–352.

22. Snider, N., Omary, M. Post-translational modifications of intermediate filament proteins: mechanisms and functions. Nat Rev Mol Cell Biol 15, 163–177 (2014).

23. Zhu, QS., Rosenblatt, K., Huang, KL. et al. Vimentin is a novel AKT1 target mediating motility and invasion. Oncogene 30, 457–470 (2011).

24. Duan W, Guo S, Huang HP, Tian Y, Li Z, Bi Y, Yi L, Cao M, Guo M, Li Y, Liu Y, Li C. High expression of NF-κB inducing kinase in the bulge region of hair follicle induces tumor. Immunobiology. 2023 Sep;228(5):152705.

25. Romanova LY, Mushinski JF. Central role of paxillin phosphorylation in regulation of LFA-1 integrins activity and lymphocyte migration. Cell Adh Migr. 2011 Nov-Dec;5(6):457–62.

26. Venet D, Dumont JE, Detours V. Most random gene expression signatures are significantly associated with breast cancer outcome. PLOS Comput Biol. 2011;7(10).

27. Avgerinosa KI, Spyrou N, Mantzoros CS, Dalamaga M. Obesity and cancer risk: Emerging biological mechanisms and perspectives. Metabolism Clinical and Experimental 2019; 92:121–135.

28. Calle EE, Rodriguez C, Walker-Thurmond K, Thun MJ. Overweight, obesity, and mortality from cancer in a prospectively studied cohort of U.S. adults. N Engl J Med. 2003 Apr 24;348(17):1625–38.

29. Renehan AG, Tyson M, Egger M, Heller RF, Zwahlen M. Body-mass index and incidence of cancer: a systematic review and meta-analysis of prospective observational studies. Lancet. 2008 Feb 16;371(9612):569–78.

30. Charlot M, Castro-Webb N, Bethea TN, Bertrand K, Boggs DA, Denis GV, Adams-Campbell LL, Rosenberg L, Palmer JR. Diabetes and breast cancer mortality in Black women. Cancer Causes Control. 2017 Jan;28(1):61–67.

31. Li Q, Hagberg CE, Silva Cascales H, Lang S, Hyvönen MT, Salehzadeh F, Chen P, Alexandersson I, Terezaki E, Harms MJ, Kutschke M, Arifen N, Krämer N, Aouadi M, Knibbe C, Boucher J, Thorell A, Spalding KL. Obesity and hyperinsulinemia drive adipocytes to activate a cell cycle program and senesce. Nat Med. 2021 Nov;27(11):1941–1953.

32. Ishaq A, Schröder J, Edwards N, von Zglinicki T, Saretzki G. Dietary Restriction Ameliorates Age-Related Increase in DNA Damage, Senescence and Inflammation in Mouse Adipose Tissuey. J Nutr Health Aging. 2018;22(4):555–561.

33. Kim D-S, Scherer PE. Obesity, Diabetes, and Increased Cancer Progression. Review. Diabetes Metab J. 2021;45:799–812.

34. Hermano E, Goldberg R, Rubinstein AM, Sonnenblick A, Maly B, Nahmias D, Li J-P, Bakker MAH, van der Vlag J, Vlodavsky I, Peretz T, Elkin M. Heparanase accelerates obesity-associated breast cancer progression. Cancer Res 2019; canres.4058.2019.

34. Maguire OA, Ackerman SE, Szwed SK, Maganti AV, Marchildon F, Huang X, … Cohen P. Creatine-mediated crosstalk between adipocytes and cancer cells regulates obesity-driven breast cancer. Cell metabolism 2021; 33: 499–512.

35. Nagy C, Einwallner E. Study of In Vivo Glucose Metabolism in High-fat Diet-fed Mice Using Oral Glucose Tolerance Test (OGTT) and Insulin Tolerance Test (ITT). J Vis Exp. 2018; 131: 56672.

36. Kleinert M, Clemmensen C, Hofmann SM, Moore MC, Renner S, Woods SC, Tschöp MH. Animal models of obesity and diabetes mellitus. Nature Reviews Endocrinology 2018;14: 140–162.

37. Nahmias Blank D, Hermano E, Sonnenblick A, Maimon O, Rubinstein AM, Drai E, Maly B, Vlodavsky I, Popovtzer A, Peretz T, et al. Macrophages Upregulate Estrogen Receptor Expression in the Model of Obesity-Associated Breast Carcinoma. Cells 2022: 11; 2844.

38. Moreno-Bueno G, Peinado H, Molina P. et al. The morphological and molecular features of the epithelial-to-mesenchymal transition. Nat Protoc. 2009; 4: 1591–1613.

39. Cervantes-Arias A, Pang LY, Argyle DJ. Epithelial-mesenchymal transition as a fundamental mechanism underlying the cancer phenotype. Vet Comp Oncol. 2013; 11: 169–184.

40. Parri M, Chiarugi P. Rac and Rho GTPases in cancer cell motility control. Cell Commun Signal 2010; 8: 23.

41. Ridley AJ; Rho GTPases and cell migration. J. Cell Sci. 2001; 114 : 2713–2722.

42. Wittmann T, Waterman-Storer CM. Cell motility: can Rho GTPases and microtubules point the way? J Cell Sci 2001; 114: 3795–3803.

43. Thomas RJ, Kenfield SA, Jimenez A. Exercise-induced biochemical changes and their potential influence on cancer: a scientific review. Br J Sports Med. 2017 Apr;51(8):640–644.

44. Mlynska A, Dobrovolskiene N, Suveizde K, Lukaseviciute G, Sagini K, Gracia BM, Romero S, Llorente A, Line A, Butkute A, Gudaite B, Venckunas T, Matuseviciene N, Pasukoniene V. Exercise-induced extracellular vesicles delay tumor development by igniting inflammation in an immunologically cold triple-negative breast cancer mouse model. J Sport Health Sci. 2025 Apr 16:101041.

45. Puurand M, Llorente A, Linē A, Kaambre T. Exercise-induced extracellular vesicles in reprogramming energy metabolism in cancer. Front Oncol. 2025 Jan 6;14:1480074.

46. Gourgue F, Mignion L, Van Hul M, et al. Obesity and triple-negative-breast-cancer: Is apelin a new key target? J Cell Mol Med. 2020; 24: 10233–10244.

47. Yelek C, Mignion L, Paquot A, Bouzin C, Corbet C, Muccioli GG, Cani PD, Jordan BF. Tumor Metabolism Is Affected by Obesity in Preclinical Models of Triple-Negative Breast Cancer. Cancers 2022, 14, 562.

48. Venneti S, Thompson CB. Metabolic Reprogramming in Brain Tumors. Review. Annual Review of Pathology: Mechanisms of Disease 2017: 12.

49. Gouirand V, Guillaumond F, Vasseur S. Influence of the Tumor Microenvironment on Cancer Cells Metabolic Reprogramming. Front. Oncol. 2018; 8:117.

50. Yang Q, Zhuo Z, Qiu X. et al. Adverse clinical outcomes and immunosuppressive microenvironment of RHO-GTPase activation pattern in hepatocellular carcinoma. J Transl Med 2024;22:122.

51. Møller LLV, Klip A, Sylow L. Rho GTPases-Emerging Regulators of Glucose Homeostasis and Metabolic Health. Cells 2019; 8: 434.

52. Machin PA, Tsonou E, Hornigold DC, Welch HCE. Rho Family GTPases and Rho GEFs in Glucose Homeostasis. Cells 2021; 10: 915.

53. Shutes A, Onesto C, Picard V, Leblond B, Schweighoffer F, Der CJ. Specificity and Mechanism of Action of EHT 1864, a Novel Small Molecule Inhibitor of Rac Family Small GTPases. Mechanisms of Signal Transduction. Journal of Biological Chemistry 2007; 282: 35666–35678.

54. Sauzeau V, Beignet J, Vergoten G, Bailly C. Overexpressed or hyperactivated Rac1 as a target to treat hepatocellular carcinoma. Pharmacological Research 2022; 179: 106220.

55. Humphries B, Wang Z, Yang C. Rho GTPases: Big Players in Breast Cancer Initiation, Metastasis and Therapeutic Responses. Cells 2020; 9(10):2167.

56. ClinicalTrials.gov. (2024). A Phase 1 open-label, first-in-human trial of oral MBQ-167 as a single agent in participants with advanced breast cancer (Identifier NCT06075810). ClinicalTrials.gov.

57. Clement E, Lazar I, Attané C, Carrié L, Dauvillier S, Ducoux-Petit M, Esteve D, Menneteau T, Moutahir M, Le Gonidec S, Dalle S, Valet P, Burlet-Schiltz O, Muller C, Nieto L. Adipocyte extracellular vesicles carry enzymes and fatty acids that stimulate mitochondrial metabolism and remodeling in tumor cells. EMBO J. 2020 Feb 3;39(3):e102525.

58. La Camera G, Gelsomino L, Malivindi R, Barone I, Panza S, De Rose D, Giordano F, D’Esposito V, Formisano P, Bonofiglio D, Andò S, Giordano C, Catalano S. Adipocyte-derived extracellular vesicles promote breast cancer cell malignancy through HIF-1α activity. Cancer Lett. 2021 Aug 21;521:155–168.

59. Barone I, Gelsomino L, Accattatis FM, Giordano F, Gyorffy B, Panza S, Giuliano M, Veneziani BM, Arpino G, De Angelis C, De Placido P, Bonofiglio D, Andò S, Giordano C, Catalano S. Analysis of circulating extracellular vesicle derived microRNAs in breast cancer patients with obesity: a potential role for Let-7a. J Transl Med. 2023 Mar 31;21(1):232.

