## Supplementary Data for "Plasma Exosomes in Insulin Resistant Obesity Exacerbate Progression of Triple Negative Breast Cancer"

**Supplementary Information**

**Supplementary Data and Figures**

In addition to the main findings presented in this manuscript, further data is provided in supplementary materials.

**Supplementary Table T1**: Combined_HFD_LFD_Mouse_GOBP_AllPathways. This table presents results from the Gene Ontology Biological Process pathway enrichment analysis, using the Fast Gene Set Enrichment Analysis method. It combines data from HFD and LFD exosome-treated mice, highlighting all enriched pathways.

**Supplementary Table T2**: Prot_Phos_Summary. This table presents a comprehensive summary of the proteomics and phosphoproteomics analysis, detailing the normalized Log_2_ intensity protein states observed under both the HFD and LFD experimental conditions.

**Supplementary Table T3**: Effects of HFD and LFD on Colony Formation in Brain and Lung Organs. Colony number, colony area and colony count of loci that expanded *ex vivo* from metastatic lesions in brain and lung are quantified, where diet of the plasma exosome donor was the independent variable, with five replicates.

**Supplementary Figures**

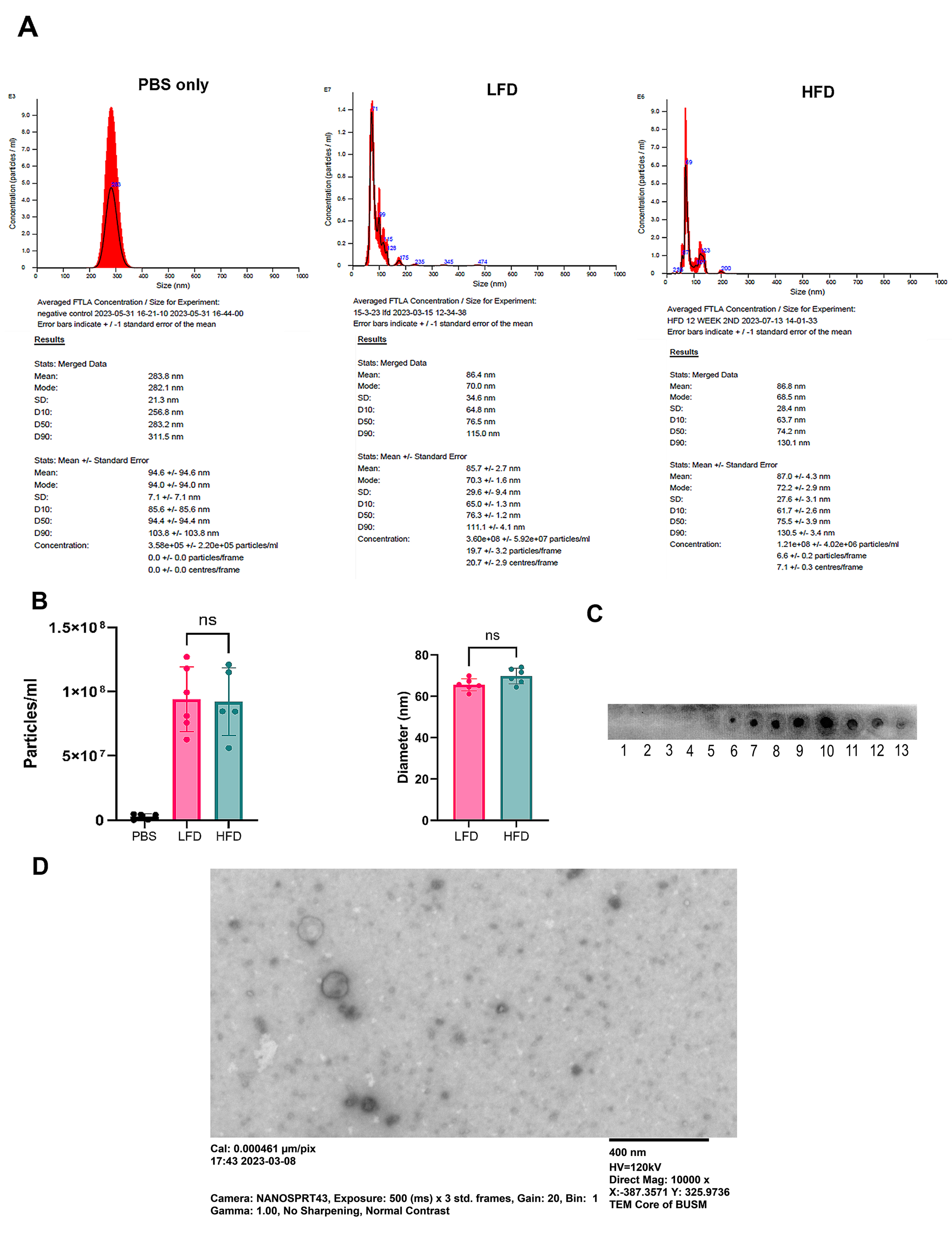

**Supplementary Figure S1:** **Plasma-derived exosome profiling control experiments using NanoSight NS300.**

(A) NanoSight NS300 size fractionation of plasma-origin exosomes. Total exosomes were purified from blood plasma of HFD and LFD fed mouse, resuspended in phosphate buffered saline pH 7.4 (PBS) and then analyzed by NanoSight NS300 to determine concentration and particle sizes. (B) Exosomes were analyzed quantitative and qualitatively, to determine their size distribution and abundance. Abundance was measured as particles/mL and mode was considered as the size distribution for each sample. Filtered PBS vehicle was used as a negative control for particle abundance. (C) CD63 dot blot confirming exosomal marker expression in pooled fractions 7–11 collected from qEV column eluates. Pooled exosomes were spotted on nitrocellulose membrane and probed for CD63 expression by immunoblotting. (D) Representative transmission electron microscopy (TEM) image of negatively stained plasma-derived exosomes. Vesicles exhibit the expected spherical morphology and size range consistent with exosomes (scale bar = 400 nm). Data are mean ± SD from N=6 independent experiments. Where ns not significant by unpaired, two-tailed t-test compared to LFD control.

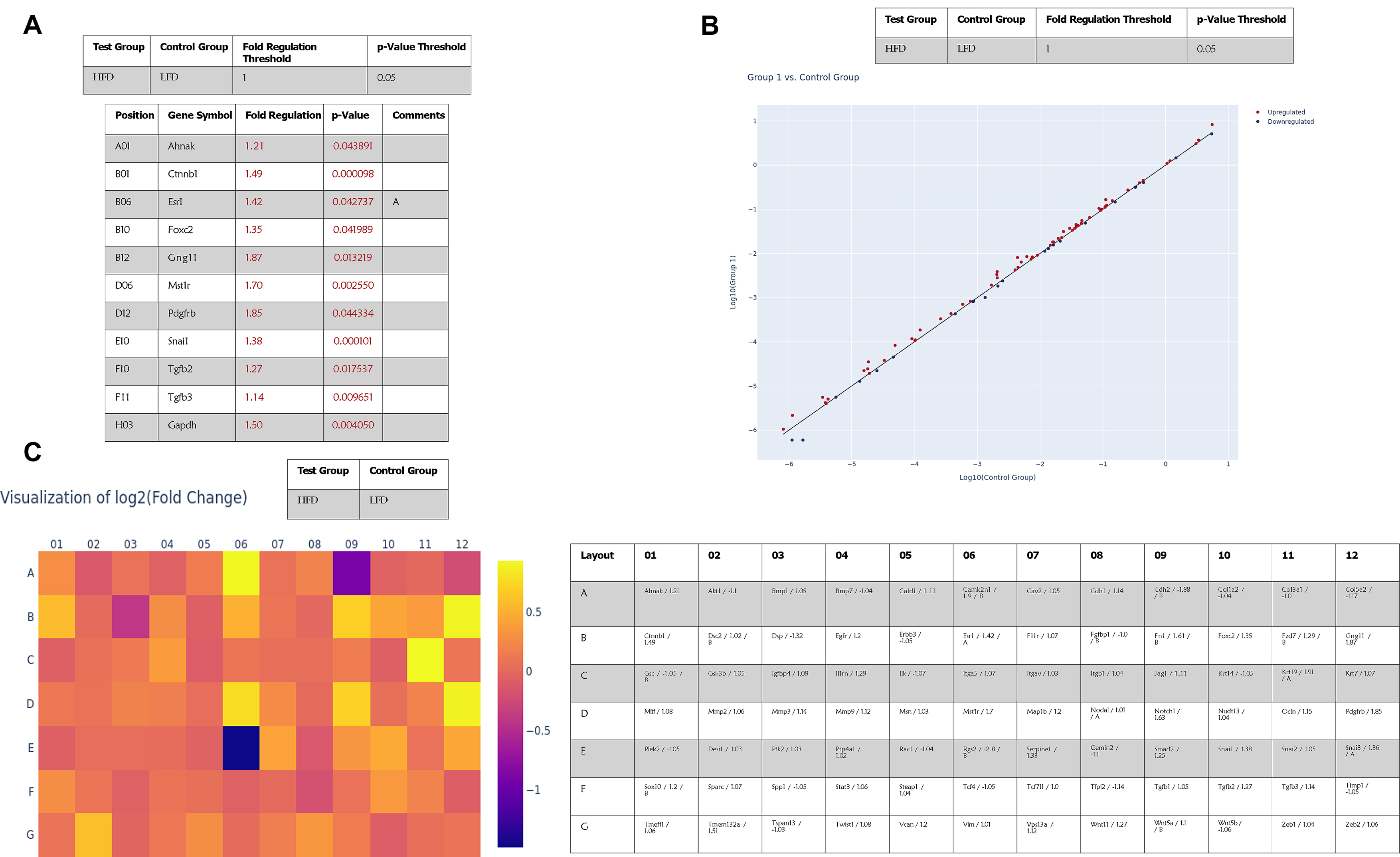

**Supplementary Figure S2: EMT array gene expression and upregulated pathways in HFD adipocyte-derived exosomes.**

(A) Fold regulation gene expression of E0771 significantly upregulated genes in HFD *vs* LFD adipocyte derived exosomes. (B) Scatter plot of HFD *vs* LFD adipocyte-derived exosomes. (C) Visualization of log2 (fold change) gene expression as a heatmap comparing HFD *vs* LFD. (Gene expression tables and plots were generated by Qiagen RT2 profiler online software. A p-value <0.05 and fold regulation >1 was considered for gene expression analysis and significant.

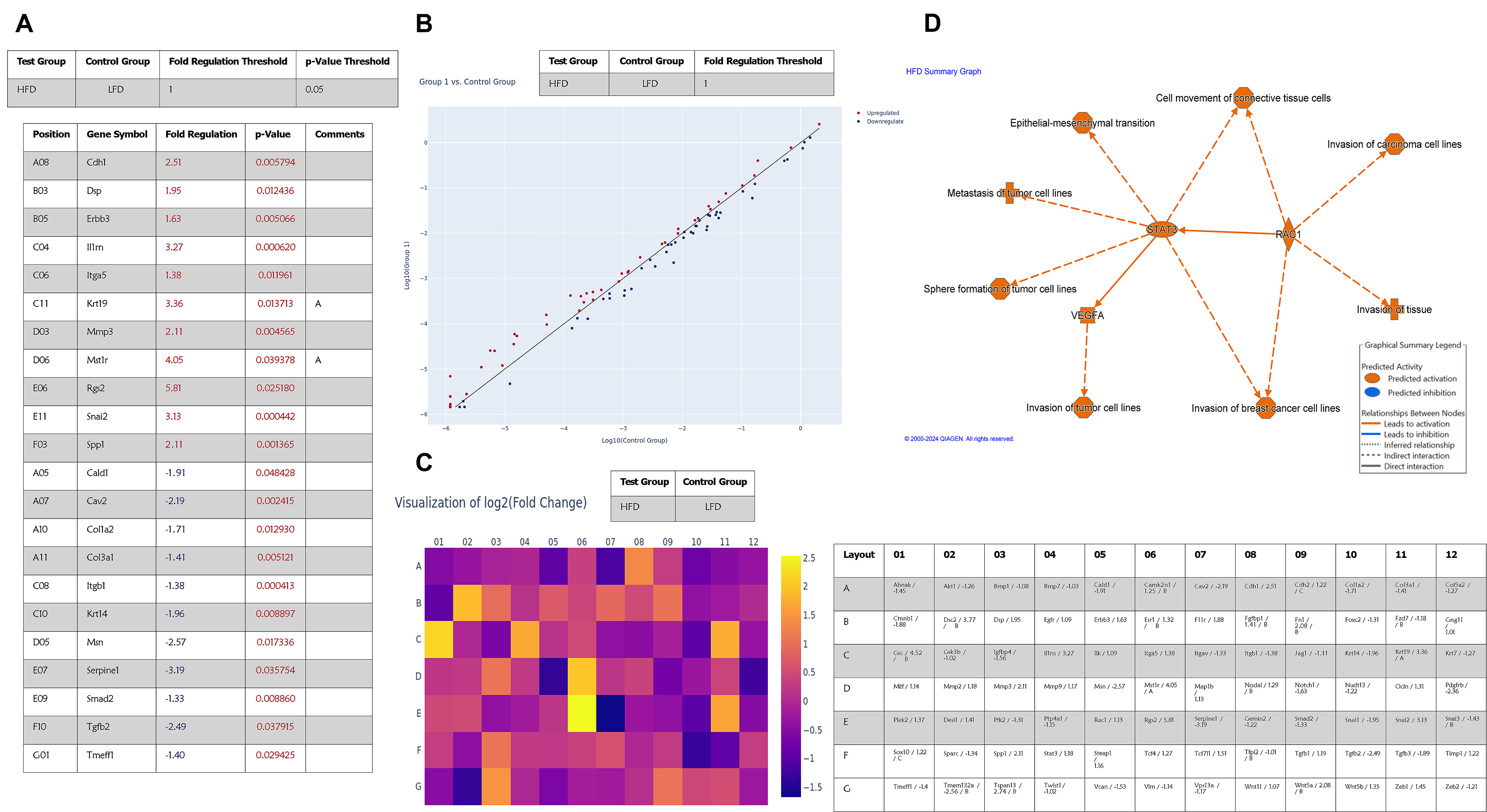

**Supplementary Figure S3: EMT array gene expression and upregulated pathways in HFD plasma-derived exosomes.**

(A) Fold regulation gene expression of E0771 significantly differentially regulated genes in HFD *vs* LFD plasma derived exosomes. (B) Scatter plot of HFD *vs* LFD plasma-derived exosomes. (C) Visualization of log2 (fold change) gene expression as a heatmap comparing HFD *vs* LFD. (D) IPA condensed graphical summary of activated pathways and functions in E0771 cells HFD-treated compared to LFD-treated. Gene expression plots were generated by Qiagen RT2 profiler online software. A p-value <0.05 and fold regulation >1 or <-1 was considered for gene expression analysis and significant. For pathway analysis, a z-score >2 or <-2 was considered as significant and included as an upregulated or downregulated pathways respectively.

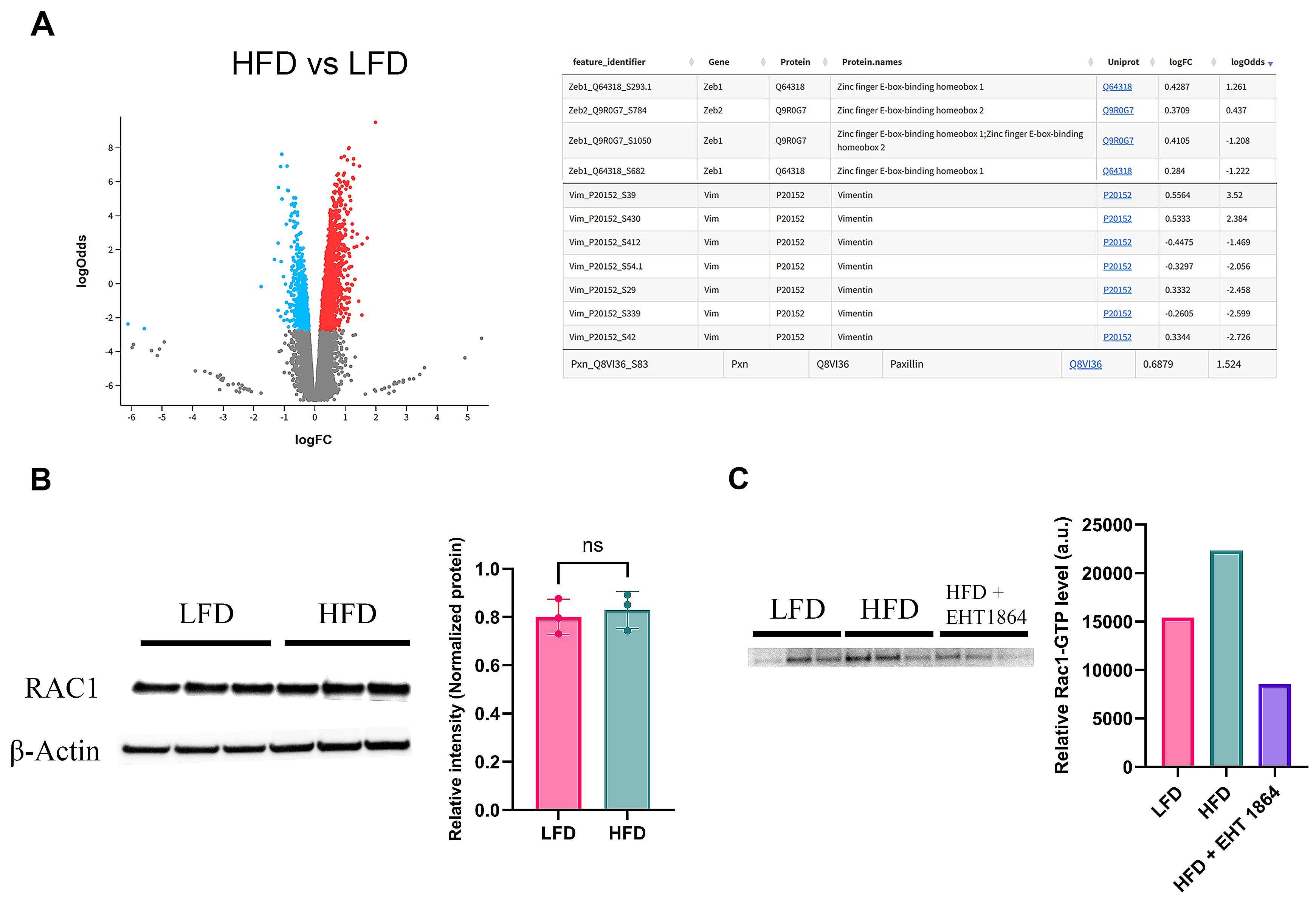

**Supplementary Figure S4: Phosphoproteomics and biochemical analyses of Rac protein signaling**

(A) Volcano plot of differentially phosphorylated proteins in E0771 cells treated with HFD- versus LFD-derived plasma exosomes. Significantly up and downregulated phosphosites (FDR < 0.05) are highlighted. Table (right) summarizes selected EMT-related targets with altered phosphorylation status, including ZEB, VIM, and PXN, with annotated phospho-sites and fold changes. (B) Western blot analysis of total Rac protein expression in E0771 cells treated with HFD or LFD exosomes. Blot quantification is shown at right. Paired t-test revealed no statistically significant difference in total Rac levels between groups. (C) GTP-bound (active) Rac1 levels were assessed by pull-down assay using the p21-binding domain (PBD) of PAK1. Western blot analysis of active Rac1 revealed a numerically higher activation in E0771 cells following treatment with HFD-derived exosomes as a confirmation assay. Data are presented as mean ± SD from N=3 independent experiments. **p < 0.01; *p < 0.001; ns, not significant by unpaired two-tailed t-test.

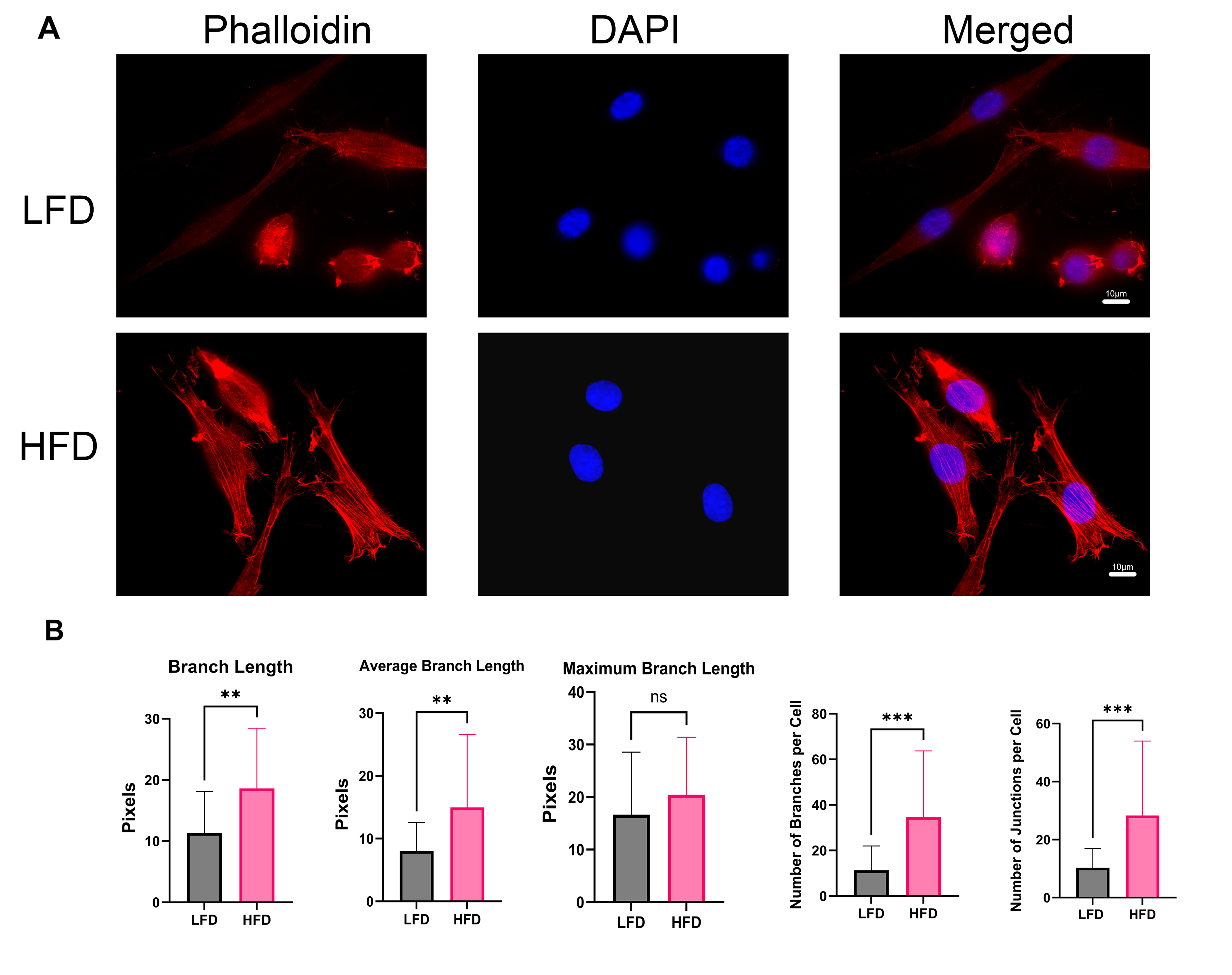

**Supplementary Figure S5: HFD-derived exosomes enhance actin stress fiber formation in E0771 cells.**

(A) Representative immunofluorescence images of E0771 cells treated with either LFD- or HFD-derived plasma exosomes. Cells were stained with Alexa Fluor™ 568 phalloidin (red) to visualize F-actin and DAPI (blue) to label nuclei. HFD-treated cells exhibit enhanced stress fiber formation compared to LFD controls. Scale bar = 10 μm. (B) Quantification of actin cytoskeletal architecture from phalloidin-stained images. Total branch length, average branch length, maximum branch length, number of branches per cell, and number of junctions per cell were analyzed using Fiji/ImageJ skeletonization and Analyze Skeleton plugin. HFD exosome treatment significantly increased total and average branch length, number of branches, and number of junctions per cell. Data are presented as mean ± SD from N=5 independent experiments. **p < 0.01; *p < 0.001; ns, not significant by unpaired two-tailed t-test.

**Supplementary Figure S6: Wound closure assay comparing the effects of EHT-1864 on HFD and LFD- plasma derived exosomes treated cells.**

Graph depicting the percentage of the wound that was closed after 24 hours across different concentrations of Rac1 inhibitor, EHT-1864 (5 µM, 10 µM, 20 µM, 25 µM, 30 µM, 40 µM, and 50 µM). (A) E0771 cells treated with LFD-plasma derived exosomes and (B) treated with HFD-plasma derived exosomes. In both HFD and LFD conditions, a dose-dependent reduction in wound closure was observed as the treatment concentration increased. HFD-treated cells had a less significant reduction in wound closure along the increasing concentrations of EHT-1864. (C) Comparison of wound closure across all concentrations in both HFD and LFD treatments. Wound closure was generally higher in LFD-treated cells at lower concentrations (5 µM and 10 µM), whereas HFD-treated cells exhibited significantly reduced closure rate at higher concentrations. (D) Morphological assessment of E0771 cells treated with HFD-derived exosomes ± EHT-1864 (25 µM) or Y-27632 (10 µM). Quantification of cell number, circularity, and perimeter is shown, along with representative phase-contrast images. Data were analyzed by two-way Anova with a Šidák’s post-hoc analysis for multiple comparison testing with statistical significance presented as: ns, not significant, *p < 0.05; **p < 0.01; ***p < 0.001; ****p < 0.0001. N=3. All data are shown as mean ± SD.

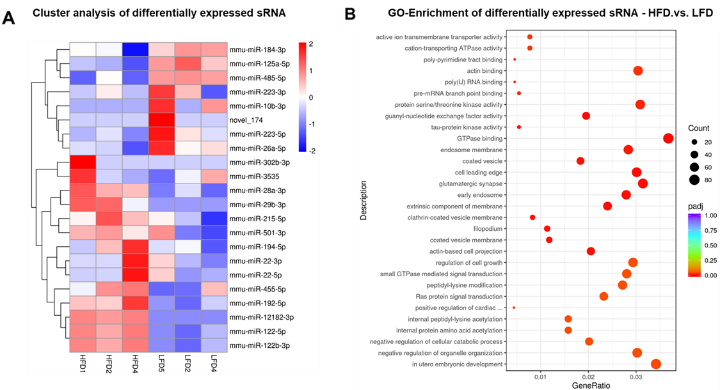

**Supplementary Figure S7: Differential expression analysis of sRNAs from plasma-derived exosomes in mice on HFD versus LFD.**

(A) Heatmap depicting differentially expressed sRNAs from triplicate samples of HFD and LFD plasma-derived exosomes in C57BL/6J mice. Z-scores for expression values are shown, representing the standardized relative expression levels across samples. (B) Gene Ontology enrichment analysis of the differentially expressed sRNAs between HFD and LFD groups, illustrating enriched biological mechanisms. GO terms are displayed with their corresponding gene ratios, adjusted p-values, and gene counts, highlighting significant pathways involved in diet-induced metabolic modulation. N=3.

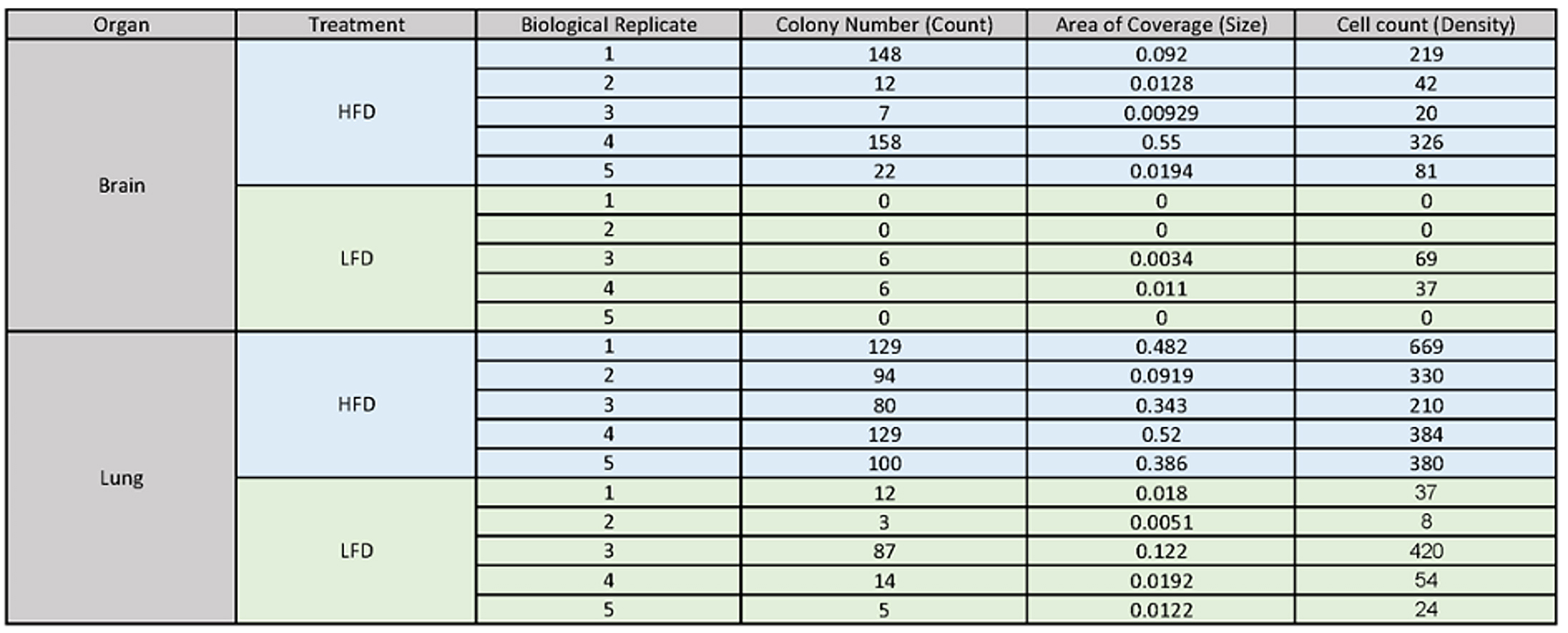

**Supplementary Table T3: Effects of HFD and LFD on Colony Formation in Brain and Lung Organs.**

Table presents data from biological replicates of brain and lung tissues obtained from C57BL/6J mice injected with E0771-GFP cells pre-treated for 72 hours with either HFD or LFD-plasma derived exosomes. Colony formation, quantified by colony number, was significantly higher in HFD-treated brain and lung samples compared to their LFD counterparts. Specifically, the brain samples from HFD-treated subjects showed large colony counts, with a peak in replicate 4 (158 colonies), while LFD-treated brain replicates either exhibited no detectable colonies (replicates 1 and 2) or very low counts (6 colonies in replicates 3 and 4). Similarly, in lung tissues, replicates treated with an HFD consistently had higher colony numbers, with replicate 1 showing the maximum count (129 colonies), compared to much lower counts in LFD-treated lung replicates, with the highest colony count being only 87 (LFD biological replicate number 3).
